# Unexpected diversity and ecological significance of uncultivable large virus-like particles in aquatic environments

**DOI:** 10.1101/2024.07.03.599014

**Authors:** Hermine Billard, Maxime Fuster, François Enault, Jean-François Carrias, Léa Fargette, Margot Carrouée, Perrine Desmares, Tom O. Delmont, Estelle Bigeard, Gwenn Tanguy, Pauline Nogaret, Anne-Claire Baudoux, Urania Christaki, Télesphore Sime-Ngando, Jonathan Colombet

## Abstract

The discovery of Jumbo phages and giant viruses of microeukaryotes has transformed our perception of the virosphere. Metagenomic and metatranscriptomic data further highlight their diversity and ecological impact. Nevertheless, sequence-based approaches fail to take into account the morphological diversity of non-cultivated viruses, resulting in our fragmented understanding of their nature and role in the environment. Here, we combined flow cytometry and electron microscopy to uncover both previously unsuspected morphological diversity as well as significant abundances of large viruses in aquatic environments. We discovered four new viral morphotypes, all of which were associated with microeukaryotes. We also obtained insights into the multi-year dynamics of the abundances of both giant microeukaryotic viruses and Jumbo phages. This work deepens our understanding of large viruses and reveals their key role as regulators of microbial communities.

## Introduction

Viruses are major actors in the environment, affecting microbial community structure and biogeochemical cycles^1–4^. Viruses forming large particles (above 0.2 µm) that infect unicellular algae^5^ or bacteria, referred to as “Jumbo” phages^6^, have been known for several decades. However, they only came to prominence following the discovery of so-called “giant” viruses (up to 1.5 µm), including the iconic mimiviruses and pandoraviruses. These giant viruses, obtained mainly by the cultivation of *Acanthamoeba polyphaga* or algal species^7–10^, have had a dramatic impact on our understanding of the evolution of viruses and blurred the boundaries between viruses and cells in terms of their physical dimensions and genomic complexity.

These culture-based studies enabled the detailed genotypic-phenotypic and biological characterization of isolated virus-host pairs^5,6,11,12^. However, the isolation of viruses is hampered by our limited ability to cultivate most microbes such as heterotrophic microeukaryotes^13^, let alone their viruses.

Metagenomics has allowed us to progressively uncover many new giant virus genomes, demonstrating that these viruses are present and diverse in most ecosystems^2,14–18^. Metagenome-assembled genomes (MAGs) can ascertain their genomic diversity, clarify their taxonomy^19,20^, and highlight their role in the evolution of cellular life forms^21–23^. For example, tens of MAGs were shown to form a diversified and prevalent group of viruses, representing a potential new phylum known as *Mirusviricota* related to the herpesviruses^24^. However, MAGs of giant viruses are often incomplete, containing little or no information about the important biological and ecological features such as the host range, mode of infection, absolute abundance, or structure and composition of the viral particle. Metatranscriptomics can provide additional information by pointing to the host of giant uncultured viruses or accessing their activity within these hosts^25^. However, as with metagenomics, this approach does not provide access to the structure and composition of viral particles or to demonstrate their absolute abundances.

To tackle the limitations of both culture and omics-based studies, Fischer et al.^26^ recently used transmission electron microscopy (TEM) to complement metagenomic approaches, thus revealing the surprising structural diversity of giant virus-like particles (VLPs) in forest soils. Although this methodology allows us to identify and categorize giant viruses based on morphological criteria, it is time-consuming and difficult to apply at high throughput. Consequently, this methodology alone is not well suited to tracking viral dynamics, thus preventing a better comprehension of the functional role played by viruses in natural ecosystems.

Flow cytometry (FC) is a rapid and inexpensive method that can be used to characterize environmental samples, leading to the absolute enumeration of nano- and microparticles. FC proved to be highly valuable for monitoring known viruses, mainly algal^27–30^, but also viruses grouped into cytometric populations characterized by specific diffusion and fluorescence signals^31–33^. However, at the current stage and in the absence of a specific marker for large viruses, the composition of these cytometric populations in environmental samples remains uncertain, even in terms of their morphology.

The recently developed FC sorting of viruses offers new perspectives to fill the gap between genotypic/morphological data and ecological significance, as it can provide absolute counts of characterized viruses after sorting. Nevertheless, to date, the application of FC sorting has remained limited to cultivable viruses^34^ or genomic characterizations^35–38^.

For the detection, morphological characterization, and ecology of large aquatic VLPs, we developed a strategy coupling TEM and FC. Applying this strategy in both directions (i.e., FC→TEM and TEM→FC) provides valuable information about large VLPs, regardless of whether they are labeled with a specific dye.

These methodological developments allowed us (i) to characterize a considerable diversity of large VLPs in three French lakes, including four new types that probably infect unicellular eukaryotes and (ii) to provide insights into the dynamics and ecology of their populations.

## Methods

### Study sites and sample collection

Samples were collected at the surface (0-40 cm) of three artificial freshwater lakes: Fargette (45°44’39’’N; 3°27’21’’E; 465 m altitude; surface area 1.2 ha; maximum depth 2.5 m), Saint Gervais d’Auvergne (SG) (46°02’15’’N; 2°48’43’’E; 680 m altitude; surface area 10.5 ha; maximum depth 4.5 m), and Chambon (45°50’22’’N; 3°30’17’’E; 490 m altitude; surface area 1.2 ha; maximum depth 6 m). These lakes are located within a 120 km radius in the French Massif Central. Fargette is a hyper-eutrophic lake, while SG and Chambon are eutrophic lakes with a significant human presence (leisure, fishing, swimming, etc.). At regular intervals, we sampled Lake Fargette from December 21, 2020, to January 18, 2024, Lake SG from February 13, 2020, to January 18, 2024, and Lake Chambon from March 23, 2022, to January 18, 2024, sampling a total of 86, 104, and 62 time points, respectively.

Samples for the counts and determination of virus-like particles (VLPs), prokaryotes (FC, TEM), and autotrophic/heterotrophic eukaryote communities (by light microscopy) were immediately fixed with 1% (v/v) formaldehyde and stored at 4^◦^C until analysis. Unfixed samples for the analysis of autotrophic/heterotrophic eukaryote communities by FC and for diversity analysis were transported at 4°C and treated in the 4 hours following sampling.

Marine samples were also collected from North Atlantic waters during the APERO expedition onboard Pourquoi Pas? in June and July 2023. One-liter volumes were collected from Nisking bottles at a depth of 2, 20, and 200 m at the three stations (PSS1: 48°27,167 N; 22°30,059 W, PSS2: 50°37,250 N; 19°7,098 W, PSS3: 47°49,846 N; 15°46,690 W). The water was prefiltered through 1.2 µm glass-fibers filters (GFC Whatman) and concentrated 9- to 10-fold by ultrafiltration using a 0.2 µm cartridge (PES Vivaflow, Sartorius). An 8 mL aliquot of the concentrate was fixed with EM grade glutaraldehyde (2% final concentration), flash frozen, and stored at –80°C until analysis.

### Abiotic parameter measurements

Dissolved oxygen content (mg.L^-1^) and temperature (°Celsius) were measured *in situ* with a submersible probe (ProDSS YSI, Yellow Springs, Ohio, USA).

### Biotic parameter analysis

#### Total pigment analysis (probe) and phytoplankton count (flow cytometry)

Total pigment content was measured (μg.L^-1^) using a submersible spectrofluorometric probe (BBE FluoroProbe, Moldaenke GmbH, DE) directly placed in the lake. Counts of pico- and nanophytoplankton populations (green algae, Cyanobacteria, Cryptophyta) were determined by FC using a BD LSR Fortessa X-20 (BD Sciences, San Jose, CA). Autotrophic organisms were categorized into five subpopulations (three subpopulations of green algae, Cyanobacteria, and Cryptophyta) according to their pigment content. Fluorescence signals from chlorophyll, phycoerythrin, and phycocyanin were collected using 405nm (50mW), 561nm (50mW), and 640nm (40 mW) lasers and 670/30, 586/15, and 670/14 filters, respectively. Green algae correspond to cells containing only chlorophyll. Cyanobacteria is the sum of phycoerythrin- and phycocyanin-rich cyanobacteria (here, we combined picocyanobacteria and large cyanobacteria). Finally, Cryptophyta corresponds to populations whose main pigments are phycoerythrin and chlorophyll.

#### Total (auto-/heterotrophic) microeukaryote counts (flow cytometry)

Counts of total (auto-/heterotrophic) microeukaryotes from unfixed samples were performed by FC using a BD LSR FORTESSA X-20 (BD BioSciences, San Jose, CA USA). The term “microeukaryote” used in this manuscript refers to unicellular planktonic eukaryotes detectable in FC or light microscopy, without size criteria.

First, autotrophic cells were targeted according to the autofluorescence of their chlorophyll content (405 nm, 50mW laser, and 635 longpass – 670/30 bandpass filters). At this stage, untargeted cells are considered heterotrophic. In each of these two populations (auto/heterotrophic), eukaryotic cells were pre-selected based on their granularity and dsDNA content using the Sybr Green I (SGI) dye (488 nm, 60 mW laser, and 502 longpass – 450/50 bandpass filters). Finally, eukaryotic heterotrophic cells were defined by mitochondrial membrane staining with 200µM Biotracker 405 Blue Mitochondria Dye (SCT135, Sigma-Aldrich, MERCK KGaA) (405 nm, 50 mW laser, and 450/50 bandpass filter). Total microeukaryotes are the sum of auto- and heterotrophic cells. This strategy assumes that all the cells of interest have mitochondria. All cytometric data were acquired and analyzed with BD FACSDiva 9.0 software.

#### Community composition of auto- and heterotrophic microeukaryotes (light microscopy)

The community composition of microeukaryotes, including eukaryotic algae, heterotrophic flagellates, and ciliates, was assessed by light microscopy. Subsamples of 1 to 4 ml (taken from the 1% formaldehyde-fixed samples) were inoculated in Utermöhl’s settling chambers containing 10 ml of < 2µm-distilled water to ensure a proper dispersion of the settled cells on the slide. The next day, slides were examined and counted under inverted epifluorescence microscopy (Zeiss Axiovert 200M, Carl Zeiss company, Oberkochen, Germany) from randomly selected transects. Cells were visualized at x400 magnification and pigmented taxa were distinguished by detecting the autofluorescence of chlorophyll a and phycoerythrin under blue light (450–490 nm) and green light (520–560 nm) excitation, respectively. All the taxa were identified with respect to their size, shape, and specific morphological characteristics (e.g., presence of flagella or cilia, colonial forms, autofluorescence, shell) to the lowest possible taxonomic level.

Net microeukaryote production (NMP) was quantified as the difference in total microeukaryote abundance (determined by FC or light microscopy) between time N and time N-1 divided by the elapsed time.

#### Diversity of eukaryotes (18S metabarcoding analysis)

For a selection of unfixed samples, microbial communities were collected on a 0.2-µm polycarbonate filter (Millipore) (until saturation, pressure < 25 kPa) and stored at –20°C until DNA extraction. The filters were covered with a lysing buffer (lysozyme 2 mg ml-1, SDS 0.5%, Proteinase K 100 µg mL-1, and RNase A 8.33 µg mL-1 in TE buffer pH 8) at 37°C for 90 min. A CTAB 10% / NaCl 5 M solution was added, and the samples were incubated at 65°C for 30 min. Nucleic acids were extracted with phenol–chloroform–isoamyl alcohol (25:24:1); the aqueous phase containing the nucleic acids was recovered and purified by adding chloroform-isoamyl alcohol (24:1). Nucleic acids were then precipitated with a mixture of glycogen 15 µg mL-1, sodium acetate 0.1M, and ethanol 100% overnight at –20 °C. The DNA pellet was rinsed with ethanol (70%), dried, and dissolved in the TE buffer. DNA was then purified using a commercial kit (NucleoSpin® gDNA Cleanup, Macherey-Nagel) and quantified by Qubit dsDNA HS kit.

The V4 region of the eukaryote SSU RNA gene was amplified using the TAReuk454FWD1 (5’-CCAGCASCYGCGGTAATTCC-3’) and TAReukREV3 (5’-ACTTTCGTTCTTGATYRA-3’) primers tagged with adaptors as recommended by the sequencing platform. Each polymerase chain reaction (PCR) was performed in a total volume of 50 µL containing 1 × final reaction buffer, 2 mM MgCl2, 0.2 mM dNTP, 250 µg mL-1 BSA, 0.4 µM each primer, 1.25 U GoTaq ® Flexi DNA Polymerase (PROMEGA), and 20 ng DNA template. The PCR protocol used an initial activation step at 94°C for 1 min, followed by 12 “three-step” cycles consisting of 94°C for 10 s, 53°C for 30 s, and 72°C for 30 s, followed by a further 18 “three-step” cycles consisting of 94°C for 10 s, 48°C for 30 s, and 72°C for 30 s, and a final 10-min extension at 72°C. PCR products were purified on agarose gel with Nucleospin® Gel and the PCR clean-up kit (Macherey-Nagel) and quantified using the Qubit dsDNA HS kit.

Library preparation, sequencing (on Illumina MiSeq, v3, 2*300 cycles), and metagenomic bioinformatic analysis (reference database: https://pr2-database.org/) were performed by the sequencing platform (Microsynth, Switzerland).

Quantification of labeled large dsDNA virus-like particles (**Fig. 1A**, Step 1a FC→TEM) and prokaryotes

**Fig. 1.**
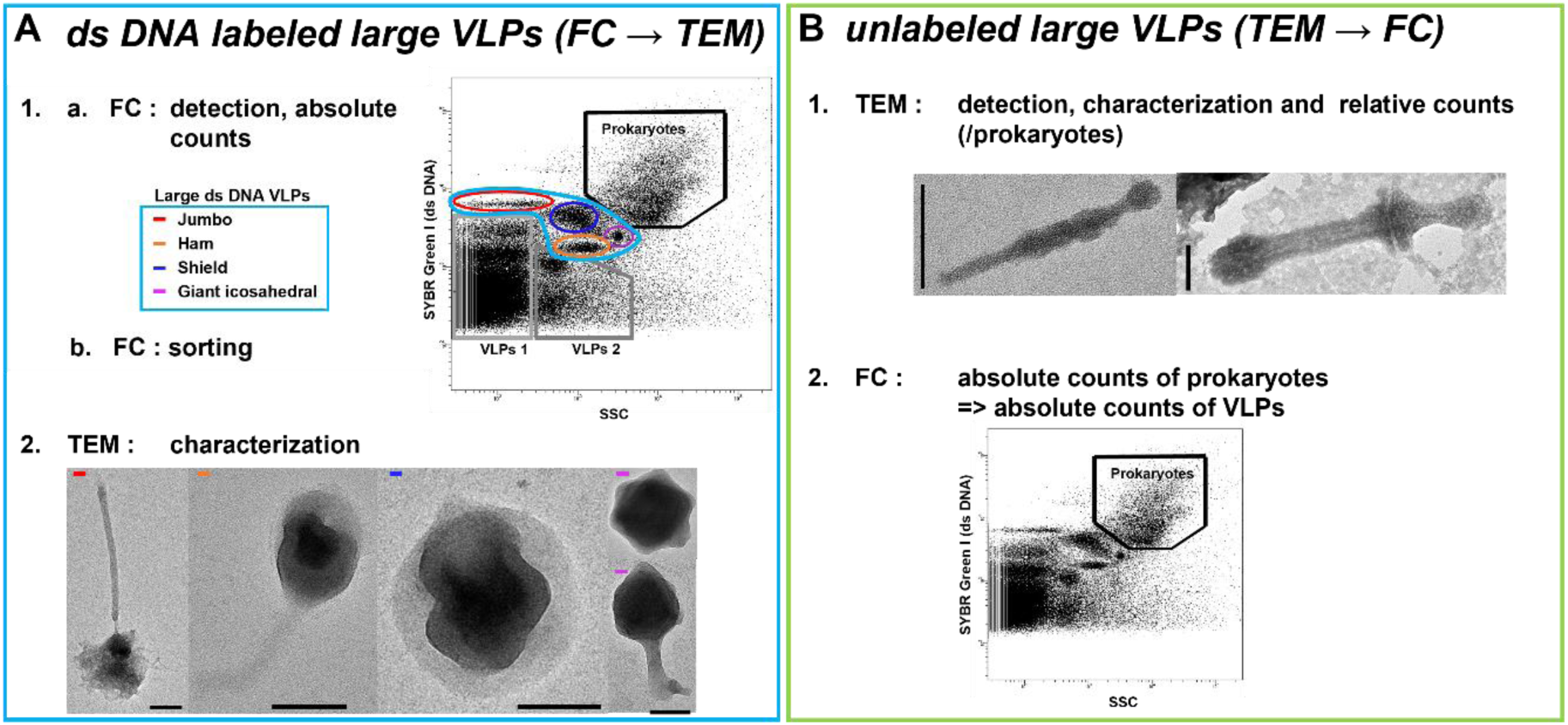
Workflow of the methodological strategy of coupling flow cytometry (FC) / transmission electron microscopy (TEM) to detect, characterize and count large virus-like particles (VLPs) ds DNA labeled (**A**) and unlabeled (**B**). **A1,** Dot plot of the gating strategy for the analysis of viral and microbial communities used in the temporal survey with emphasis on large dsDNA-labeled VLPs determined according to their ds DNA content (SYBR Green I) and side scatter (SSC) intensities. 8 populations were considered: Jumbo, Shield, Ham, giant icosahedral VLPs (the sum of these VLPs constitutes the large dsDNA VLPs), VLPs 1 and 2 and prokaryotes. **A2**, Negative staining electron micrographs micrographs of large dsDNA-specific VLPs sorted by FC into FC-identified gates. **B1**, Negative staining electron micrographs of large unlabeled specific VLPs detected in environmental samples. **B2**, Dot plot of the gating strategy for the analysis of prokaryotes used to convert relative counts of unlabeled large VLPs into absolute counts. Scale bars = 100 nm.

Counts of dsDNA VLPs and prokaryotes from fixed samples were performed in triplicates by FC using the dsDNA SYBR Green I (SGI) dye (S7585, Invitrogen, Thermo Fisher Scientific) according to Brussaard (2004)^29^ with a BD FACSAria Fusion SORP (BD Sciences, San Jose, CA, USA) equipped with an air-cooled laser delivering 50 mW at 488 nm with 502 longpass, and 530/30 bandpass filter set-up. All cytometric data were acquired and analyzed with BD FACSDiva 9.0 software.

Microbial and VLPs were divided into subpopulations according to their intensity on the side scatter (SSC) and SGI signals. First, prokaryotes were targeted. Second, two populations (VLP1 and VLP2) were characterized by low SSC and/or low SGI signals. VLP1 is dominated by small bacteriophages (< 200 nm)^29,39^, although small eukaryotic algal viruses such as small genome-sized *Heterosigma* or diatom viruses can also display similar low fluorescence signatures^40–43^. VLP2 was mainly composed of tailless icosahedra of around 100 nm in diameter (data not shown), commonly identified as dsDNA algal viruses^28,41,44^. Between these microbial and small VLP populations, four populations were identified on cytograms characterized by high SSC and/or high SGI signals. Considering the morphology of the VLPs contained therein (observed and regularly verified after FC sorting by TEM), these four populations were named Jumbo (Jumbo-like phage), Shield, Ham, and giant icosahedral VLPs (GIV).

Jumbo, Shield, Ham, and GIV form a group of large dsDNA VLPs.

The gating and counting strategy are provided in **Fig. 1A**. Dates with gate overlaps, notably from prokaryotes or debris populations, were excluded from the analysis.

Note that we were unable to identify cytometric populations corresponding to Snake and Sword VLPs on cytograms after FC sorting. For these populations, only TEM→FC counts were available (**Fig. 1B**).

Fluorescence-activated large virus-like particle sorting (**Fig. 1A**, Step 1b FC→TEM)

For VLP sorting, we selected dates with remarkable virus populations on account of their fluorescence signals and intensity: Fargette on March 1, 2022, SG on February 14, 2022, and Chambon on December 16, 2022, and January 12, 2024.

VLP sorting (1% formaldehyde-fixed sample) was performed with the FACSAria Fusion SORP (BD Biosciences, San Jose, CA, USA) flow cytometer using a 488 nm argon laser for excitation. The “single cell” sort mode was used to ensure that the sorted drops were free of contaminating particles and that the target particles were centered within the deflected drop.

Sorting instruments and reagents were decontaminated as recommended by the manufacturer (Prepare for Aseptic Sort, BD Biosciences, San Jose, CA, USA). The threshold of the cytometer was triggered at the minimum in the green fluorescence channel, and the sorting gates were based on SGI fluorescence and SSC.

From each of the distinct populations mentioned above, between 3.10^5^ and 3.10^6^ VLPs were sorted for TEM identification. For this sorting, we used sterile NaCL (2.24g.L^-1^) diluted in ultrapure water solution as sheath fluid. This NaCL concentration was determined to permit correct deflection (concentration range test, data not shown) in order to prevent salt contamination of the TEM observations.

*Quantification of specific large virus-like particles with or without flow cytometry signal* (**Fig. 1B**, Step 1 and 2 TEM→FC) *and morphological characterization* (**Fig. 1A** Step 2, **1B** Step 1)

VLPs and microbial communities in 1% (v/v) formaldehyde fixed samples were collected by centrifugation at 20,000 x *g* for 20 min at 14°C directly on 400-mesh electron microscopy copper grids covered with carbon-coated Formvar film (AO3X, Pelanne Instruments, Toulouse, France). Particles were over-contrasted using uranyl salts. Specific viruses were detected, characterized, and counted by TEM using a Jeol JEM 2100-Plus microscope (JEOL, Akishima, Tokyo, Japan) equipped with a Gatan Rio 9 CMOS camera (Gatan Inc., Pleasanton, USA) operating at 80 kV and x50,000 to x150,000 magnifications. TEM images were acquired and measurements made with DigitalMicrograph-GMS 3 (Gatan Inc., Pleasanton, USA). Resizing as well as light and contrast corrections were carried out with ImageJ^45^ or Photos Microsoft (Microsoft Corporation, Redmond, Washington, USA). Scale bars were retraced and formatted manually.

The number of specific VLPs with (Ham, Shield) or without FC signals (Snake and Sword) resulted from the multiplication of the VLP/prokaryote ratio determined by TEM by the prokaryote concentration obtained by FC (**Fig 1B***, Step 2 TEM→FC*).

The terms “large,” “Jumbo,” or “giant” were used in relation to the size of VLPs (length >200 nm on one of their axes) and FC characteristics (high values of SGI (dsDNA) or SSC (complexity/size) mean fluorescence intensities).

For VLPs presenting an identified cytometric population (Shield, Ham), we validated the reciprocity between the methods FC→TEM and TEM→FC by their positive correlation (**Extended Data** Fig. 1). In the analysis of their dynamics, we present the results of the method with the maximum number of points (i.e., FC→TEM for Shield VLPs and TEM→FC for Ham VLPs). Giant icosahedral VLPs were only counted by FC. For VLPs with no detectable cytometric signature using SGI (Snake and Sword), the TEM→FC approach was the only methodology permitting the detection, identification, and quantification of VLPs in this study.

Large dsDNA VLP populations (Shield, Ham, Giant Icosahedral, Jumbo-like phages, Snake and Sword VLPs) were combined to consider the total community of large VLPs. Jumbo-like phages were removed to consider the potential population of large microeukaryote VLPs.

Net viral production was quantified for each large VLP and for the sum of large microeukaryote VLPs as the difference in VLP abundance between time N and time N-1 divided by time. A theoretical viral death rate of microeukaryotes was evaluated as the net viral production divided by the burst size (BS). BS was determined from TEM observations of host lytic events for Shield, Ham, Snake, and Sword VLPs. For GIV, we used a theoretical BS of 50. For the total large microeukaryote VLPs, we used the average of the BS observed in TEM and in theory for each of the quantified viral types. We considered the estimate of the viral death rate of microeukaryotes to be minimal, as it does not take into account the viral decay, which was not measured here (**Table 1A**).

**Table 1.**
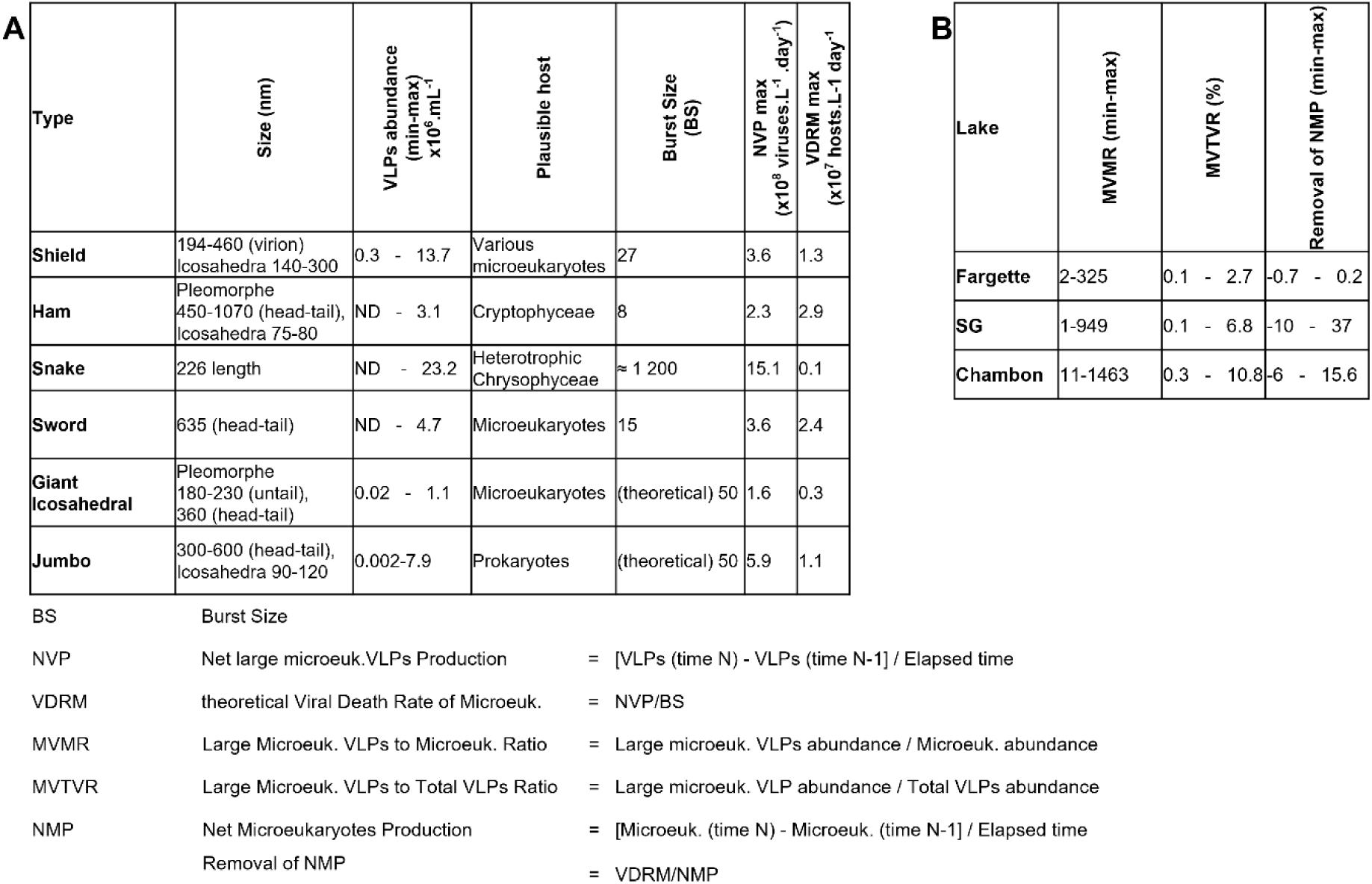
Morphological and ecological characteristics of the viruses-like particles considered in this study. Burst size was counted from transmission electron microscopy observations of host lytic events. ND: not detected.

We estimated the removal of NMP by large microeukaryote VLPs by applying the ratio between the viral death rate of microeukaryotes and NMP. It was only estimated at the dates for which autotrophic and heterotrophic microeukaryote counts by light microscopy or mitochondrial tagging were available (**Table 1B**).

## Data analysis

All statistical analyses were performed using R software (version 4.1.3, R Foundation for Statistical Computing, Vienna, Austria). Potential relationships among all variables were tested by pairwise correlations (Spearman correlation analyses).

## Results and discussion

### Large VLPs: Abundant and dynamic players in viral ecology

Data from cultures^5,6,10–12^, microscopic observations^26^, and genomic analyses^2,14–17^ have led to fundamental discoveries in the study of large viruses. However, in the absence of a methodology to detect, characterize, and quantify their non-cultivable representatives, their diversity and ecological importance remains underexplored. To fill this knowledge gap, we coupled FC and TEM (**Fig. 1**) to study large viral particles in aquatic ecosystems. FC not only allowed the detection and absolute counts of large VLPs^29^ carrying dsDNA genomes

fluorescently labeled with SYBR Green I (SGI) but also enabled the comparison of corresponding subpopulations on the basis of fluorescence signals associated with SSC and dsDNA fluorescence^28,31,34^ (**Fig. 1A**). TEM allowed us to assess the homogeneity and morphological characteristics of the populations sorted using FC as well as the visualization and quantification of large VLPs (**Fig. 1A-B**) from environmental samples that were not labeled in the FC analysis.

Using this protocol, the populations of large VLPs were morphologically characterized and quantified in multiple samples collected at regular intervals over successive years (2020-2024) in three artificial eutrophic freshwater lakes located in the French Massif Central (Lakes Fargette, Saint Gervais d’Auvergne (SG), and Chambon).

Large VLPs (FC labeled or not) were consistently present in the studied systems, with abundances ranging from 0.02 to 2.8 x 10^7^ VLPs.mL^-1^, i.e., from 0.2 to 11.2% of total VLPs. Their dynamics were characterized by phases of production in spring or late autumn alternating with low detection phases (**Fig. 2**). Notably, among the large VLPs, up to 78% were unlabeled (**Fig. 2**). To decipher the significance of aquatic large VLPs, most of which did not resemble the previously characterized VLPs, we explored their morphological diversity and ecological patterns (i.e., dynamic, interactions with microbial communities) while separating the VLPs into two groups, namely Jumbo phages and large VLPs likely associated with microeukaryotes. The term “microeukaryotes” used in this manuscript refers to unicellular planktonic eukaryotes detectable in FC or light microscopy, irrespective of their size.

**Fig. 2.**
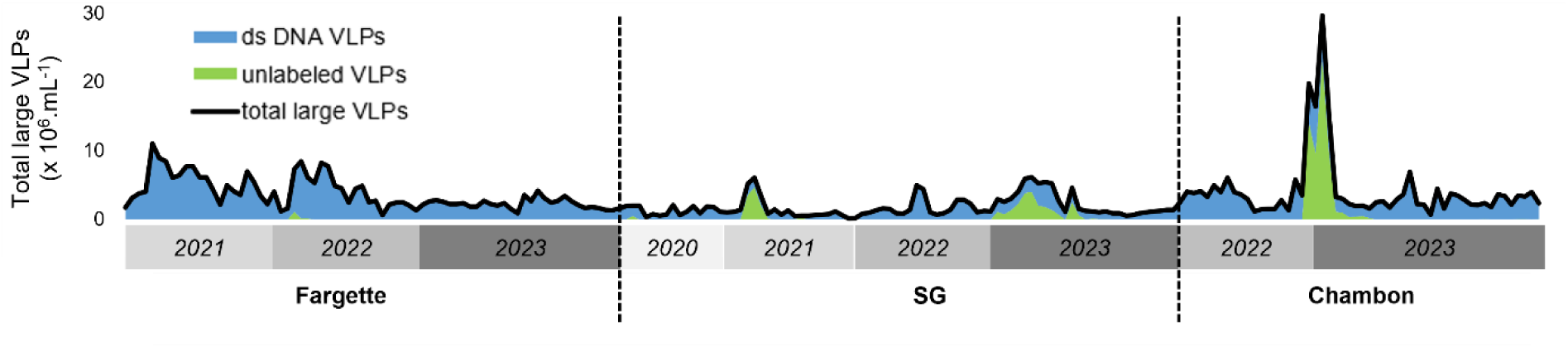
Seasonal dynamics of the abundance of total community of large virus-like particles (VLPs) (labeled and unlabeled), in lakes Fargette, SG and Chambon. Each data represents the average of triplicates, n= 209.

### Overlooked diversity and ecological significance of Jumbo phages

Although a number of Jumbo phages were previously isolated and cultivated^11,46^, their diversity and ecology in aquatic environments were almost exclusively analyzed by metagenomics^2,16^. These studies uncovered an enormous diversity, suggesting that Jumbo phages are important, yet underestimated components of microbial communities and food webs^2^. To examine this viral component, we identified a specific cytometric population that corresponds to Jumbo-like phages, as confirmed by TEM observations following FC sorting (**Fig. 3A-B**). The population, named Jumbo, was characterized by a low SSC and a high level of fluorescence, which is a proxy for the large genome sizes of Jumbo phages (> 200 kb) and consistent with what was previously demonstrated for the iconic Jumbo coliphage T4^29^. TEM showed capsid sizes between 90 and 110 nm in diameter, with total lengths between 270 and 467 nm (**Fig. 3B-C**). The overall organization of VLPs was characterized by head-tailed phages of the class *Caudoviricetes*^11^. The abundances of the Jumbo VLPs reached 7.9 x 10^6^ VLPs.mL^-1^, and their dynamics showed irregular phases of development (**Fig. 3D**). For example, there was a noticeable peak in Lake SG from March 8 to July 4, 2022. Jumbo VLPs were detected in all the years under investigation. As expected, this population showed a positive correlation with prokaryotes (R² = 0.63, p-value < 0.00001).

**Fig. 3.**
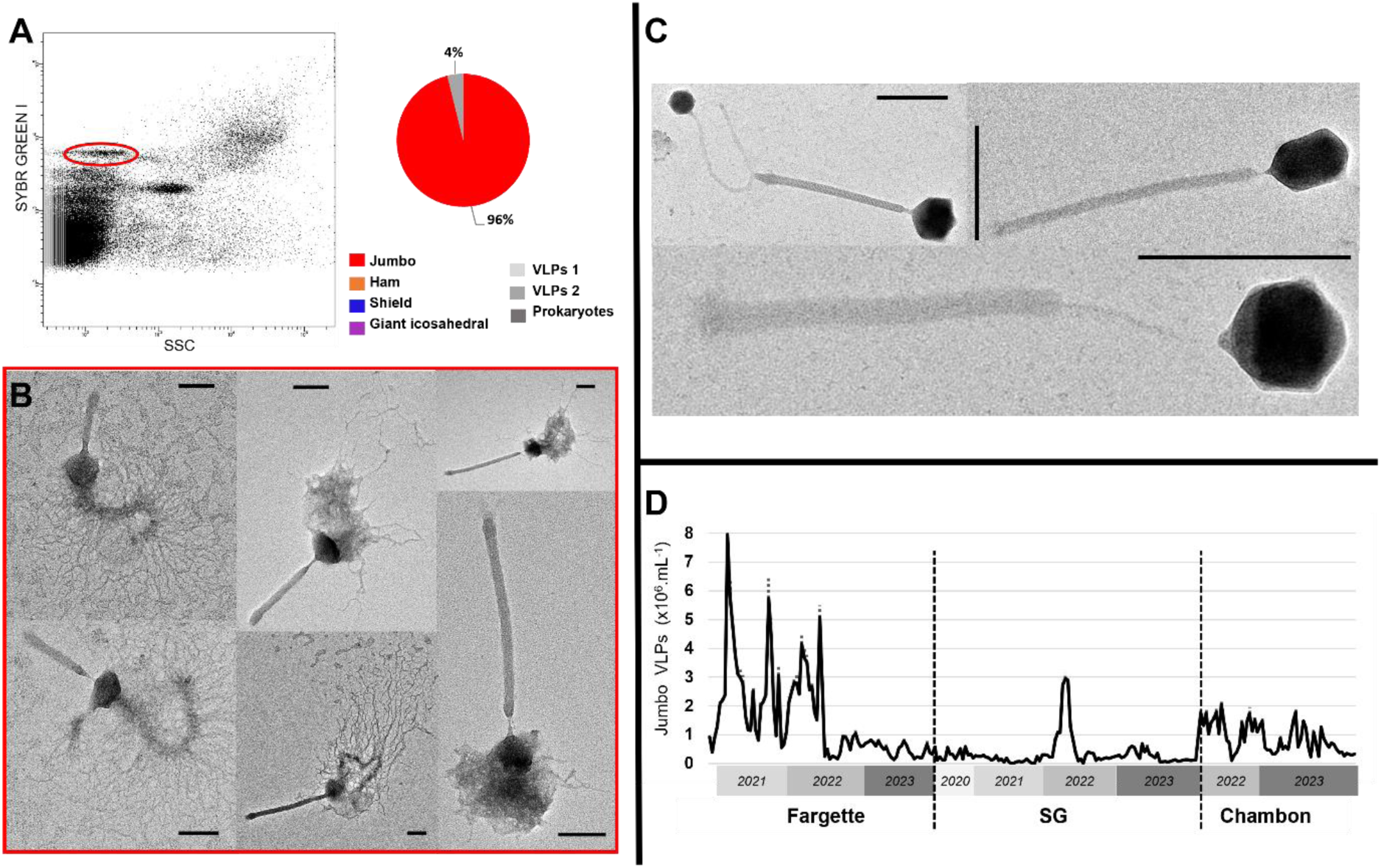
Detection, morphological and ecological characterization of Jumbo-like phage populations identified in eutrophic French lakes. **A,** Flow cytometry (FC) detection of a Jumbo- like phage population and diversity of entities recorded by transmission electron microscopy of this sorted population. **B,** Negative staining electron micrographs displayed Jumbo-like phages obtained in the sorted population. Note the deleterious effect of sorting with visible damage on the capsid of Jumbo-like phages on which the molecular structure comes out of the head. **C**, Negative staining electron micrographs of Jumbo-like phages detected in environmental samples. Scale bars in B = 100 nm, C = 200 nm. **D**, Seasonal dynamics of Jumbo-like phage abundances, in lakes Fargette, SG and Chambon. Each data represents the average of triplicates, dotted lines indicate standard deviation. n = 252.

In the absence of evident burst events, we used a theoretical BS of 50, within the range of a previous report^47^, to estimate the high maximum potential mortality of 1.1 x 10^7^ hosts.L^-1^.day^-1^ (**Table 1A**).

The potential Jumbo phages observed in the raw environmental samples (**Fig. 3C and Extended Data** Fig. 2) were more morphologically diverse than those belonging to the Jumbo population sorted by our defined cytometric gate. Indeed, whereas the VLPs of the Jumbo FC population were fairly homogeneous in shape and size, environmental Jumbo-like VLPs tended to be more heterogeneous. Although some particles consisted of a well-defined capsid and tail, others had tails that were covered by complex sheath-like structures or that were remarkably long, up to 2,200 nm in length. Even though we cannot exclude the possibility that all these diverse tailed particles were in our defined Jumbo cytometric gate, most were likely rare and outside any cytometric gate.

These results show that Jumbo-like phages are neglected in the aquatic environment in terms of their diversity, abundance, and induced mortality. Indeed, that they could be of major importance in the microbial loop^2^.

### Large VLPs of microeukaryotes: Amazing diversity and unexpected ecological significance

As in the case of Jumbo phages, the diversity and ecology of large aquatic VLPs of microeukaryotes remains largely underexplored. Here, we discovered four new abundant large viral morphotypes. Whereas VLPs resembling the Shield form were previously detected in soil samples, dubbed as the “Christmas star”^26^, the Ham, Snake and Sword VLPs present completely new shapes never before observed. In addition, we provide the morphological characterization and monitoring of abundant large icosahedral VLPs and highlight the ecological significance of the total community of large VLPs of microeukaryotes.

#### Ham virus-like particles: Atypical morphology of a new viral type

During our study, we detected a particular cytometric signature corresponding to a new type of VLPs, which we named Ham. The average fluorescence intensities derived from the dsDNA labeling and SSC of these particles were intermediate, being situated between VLPs2 and prokaryotes, suggesting that they have large genomes (**Fig. 1A, 4A**).

The Ham VLP is a polymorphic particle with lengths ranging from 450 to 1070 nm (**Fig. 4**). It is characterized by an ovoid head ranging from 240 to 500 nm in length and from 135 to 190 nm in width, a tail surrounded by an apparently helical assembly ranging from 140 to 550 nm in length and from 40 to 50 nm in width. The end of the tail is decorated by fibers. Some morphotypes have a bulge at the end of the tail. A notable feature is the presence of an icosahedral structure of 75-80 nm in diameter inside the head. The reproducibility of the observations of this icosahedral structure inside the putative virion suggests that this is an integral part of the Ham VLP. Two or three icosahedrons inside the largest form were occasionally observed (**Fig. 4B**). These multi-icosahedron forms were rare (< 1%), except in Lake Chambon from December 20, 2023, to January 10, 2024, in which they could represent up to 64% of the total Ham VLPs counted. To our knowledge, this is the first description of a head-tail VLP with an icosahedral structure covered by an external layer. Morphologically, the closest relative appears to be the enigmatic virions of Meelsvirus, which have head-tail particles with the head containing an ovoid nucleocapsid^48^.

**Fig. 4.**
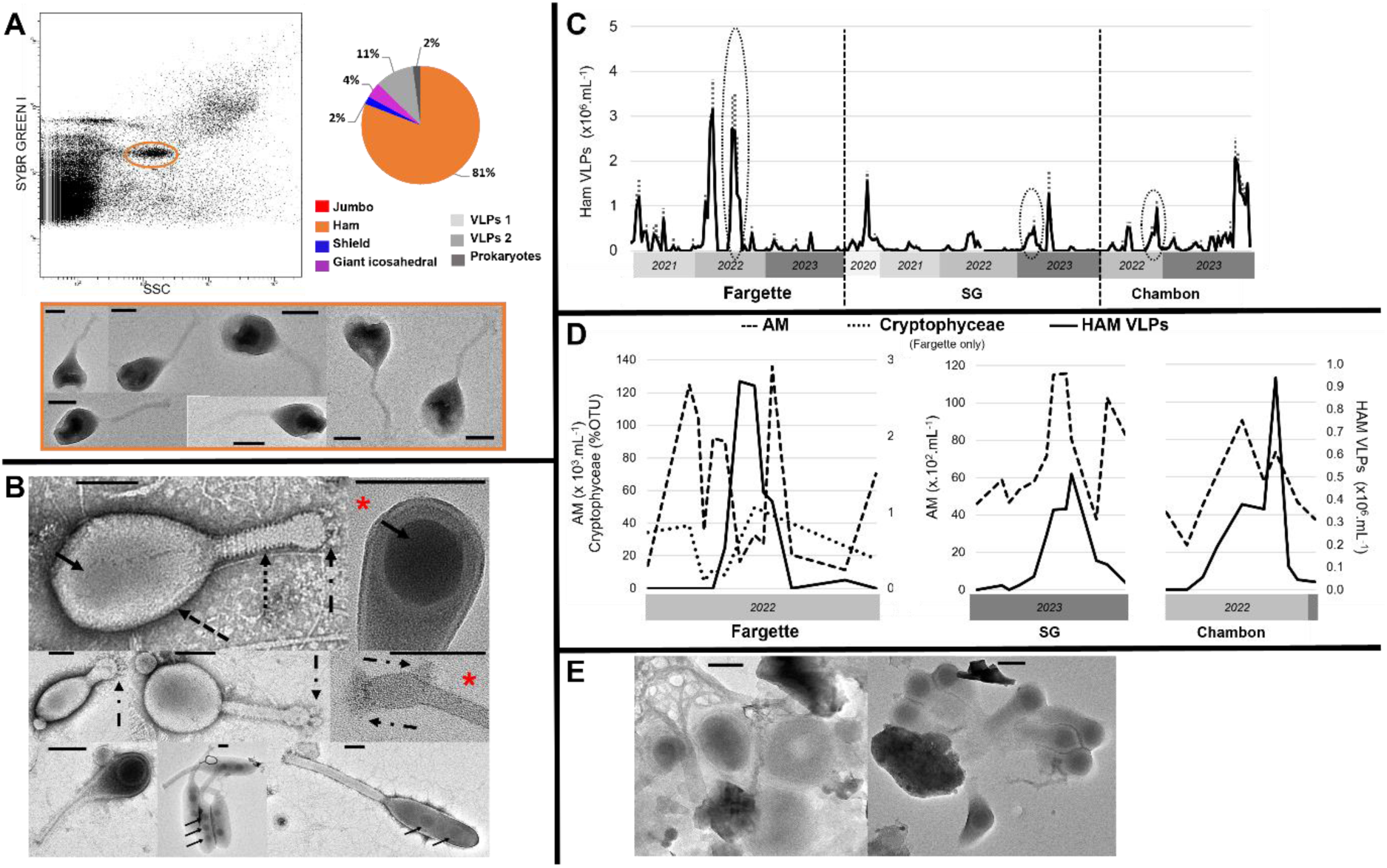
Detection, morphological and ecological characterization of Ham virus-like particles (VLPs) detected in eutrophic French lakes. **A**, Flow cytometry (FC) detection of remarkable population of Ham VLPs and diversity of entities recorded by transmission electron microscopy after FC sorting in the corresponding population with micrographs of the Ham VLPs sorted. Note the deleterious effect of sorting with visible damage on the capsid of Ham VLPs. **B,** Negative staining electron micrographs of Ham VLPs in which we detected an, two or three icosahedral structures (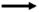) contained within a surrounding structure with a head (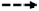)-tail (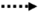) morphology. **\*** Illustrated zoom on head or tail. Note the spiral-shaped molecular structure around the tail (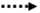) and the presence of fibers (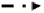) at the end of the tail. **C**, Seasonal dynamics of the abundance of Ham VLPs, in lakes Fargette, SG and Chambon. Each data represents the average of triplicates, dotted lines indicate standard deviation. n=252. **D**, Focus on remarkable infection periods on the covariations of Ham VLPs and autotrophic microeukaryotes (AM) and relative abundance of Cryptophycaea class (% OTU) in lake Fargette from March 29, 2022 to August 30, 2022, in lake SG from January 16, 2023 to April 12, 2023, and in lake Chambon from September 23, 2022 to January 3, 2023. **E,** Negative staining electron micrographs of Ham VLPs derived from a lytic event. Scale bars A, B, E = 100 nm.

Ham VLPs were encountered in all three lakes considered in this study and reached 3.1 x 10^6^ VLPs.mL^-1^ with a sudden and massive development strategy (**Fig. 4C**). The proliferation dynamics of Ham VLPs strongly correlated with the FC phytoplanktonic microeukaryote population identified as Cryptophyta (R² = 0.3, p-value < 0.00001). Metabarcoding data showed that at the time of the VLP bloom, eukaryotic communities were dominated by Cryptophyceae, which further suggests that Ham VLPs are produced by the Cryptophyceae species (**Fig. 4D)**. Notably, the interaction between Ham VLPs and autotropic microeukaryotes was also supported by the “prey-predator” model (**Fig. 4D**). This model of interaction has been frequently observed for large algal viruses^33,49–53^. Without further precisions about the specific identity of the host, these observations suggest that Ham VLPs infect autotrophic microeukaryotes. The pleomorphism of Ham VLPs suggests that it could be represented by different phylotypes, a possibility reinforced by the ubiquity of Ham VLPs and their development at different times of the year under contrasting environmental conditions (temperature range 5 to 20 °C). However, its development strategy and malleable morphology could also support the hypothesis of specific polymorphic VLPs.

The small number of VLPs counted after the observation of cell remnants (**Fig. 4E**) suggests that the BS was reduced to a few units. We estimated this VLP to be involved in the mortality of up to 2.9×10^7^ hosts.L^-1^.day^-1^ (**Table 1A**).

#### Shield virus-like particles: A new universal viral phylum?

Like Ham VLPs, Shield VLPs exhibited a specific cytometric signature with fluorescence intensities in the range of small prokaryotes and slightly lower than Jumbo-like phages. The cytometric signatures of Ham and Shield VLPs correspond those of large genome dsDNA algal viruses such as *Phaeocystis* or *Emiliana huxleyi* viruses^27–31^ or *Bodo saltans* virus infecting a heterotrophic flagellate^54^, suggesting a large genome (**Fig. 1A, 5A**).

Shield VLPs have an electron-dense inner capsid (140-300 nm in diameter) surrounded by a less electron-dense spherical outer structure (194-460 nm in diameter) (**Fig. 5A-B, Extended data Fig. 3A**). The inner capsid features a pentagonal planar projection with a 5-9 nm thick border. Some representatives of Shield VLPs suggest that the external structure is composed of a tegument and an envelope, as is characteristic of viruses in the order *Herpesvirales* (**Fig. 5B**). However, the nature of this envelope-like structure is unclear, as it could be lipidic, as in herpesviruses, or based on glycans, as in *Megavirinae*^55,56^. Unlike *Megavirinae*, the hairy appearance resulting from the arrangement of fibrils has not been clearly observed, and some Shield representatives listed here are smaller than *Megavirinae*. However, one example of a particular *Megavirinae,* namely Cotonvirus (400 nm in capsid diameter), shows that the density of fibrils on the capsid can have a smooth appearance under TEM^57^.

**Fig. 5.**
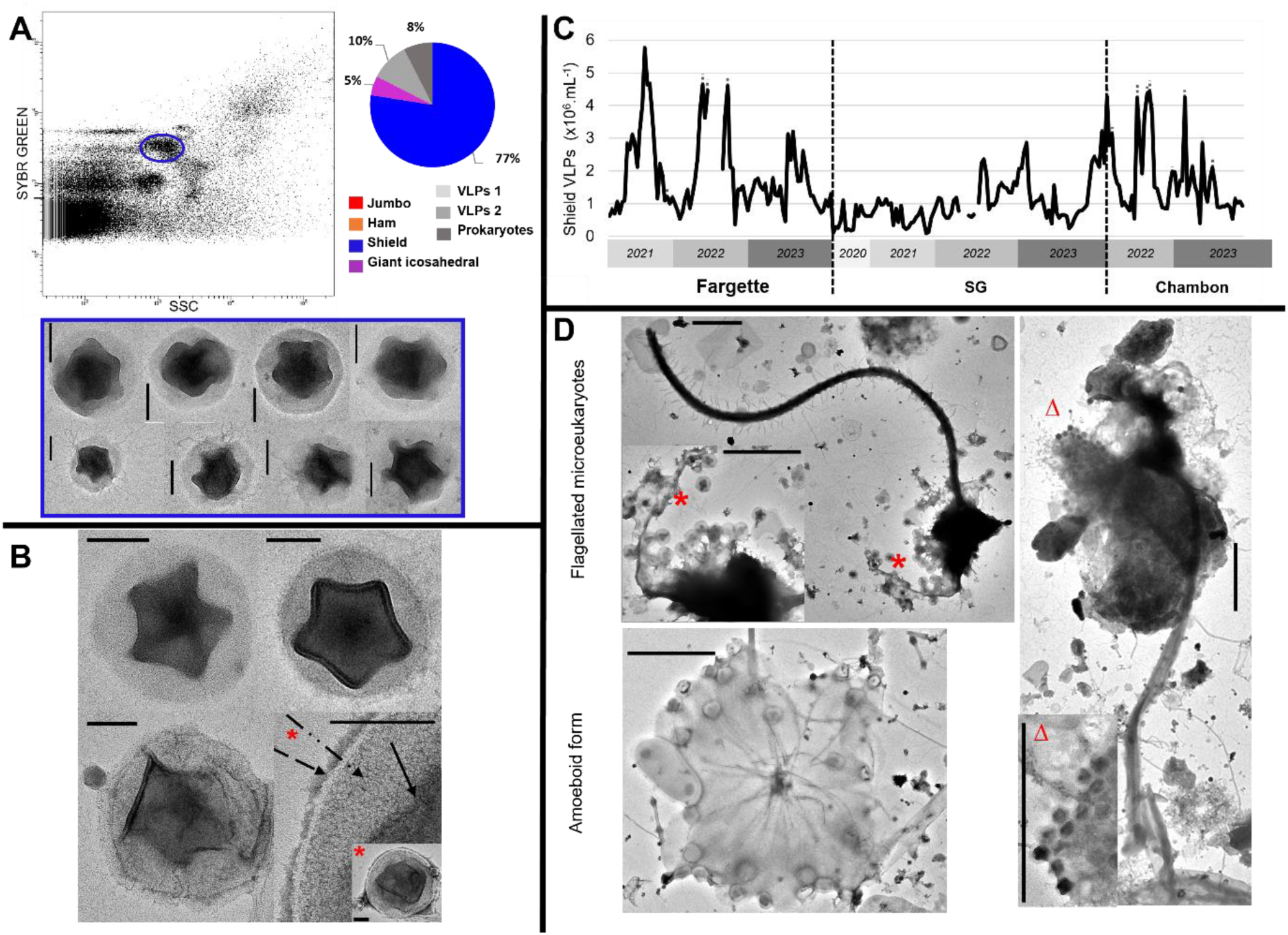
Detection, morphological and ecological characterization of Shield virus-like particle (VLPs) detected in eutrophic French lakes. **A**, Flow cytometry (FC) detection of remarkable population of Shield VLPs and diversity of entities recorded by transmission electron microscopy after FC sorting in the corresponding population with negative staining electron micrographs of the Shield VLPs sorted. **B**, Negative staining electron micrographs of Shield VLPs in which we identified an inner pentagonal structure (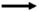) surrounded by a tegument (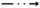) and an envelope (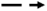)-like structure. Note the stargate-like motif in the center of the inner structure. **C,** Seasonal dynamics of the abundance of Shield VLPs, in lakes Fargette, SG and Chambon. Each data represents the average of triplicates, dotted lines indicate standard deviation. n=243. **D**, Negative staining electron micrographs of Shield VLPs derived from lytic events of flagellated microeukaryotes or amoeboid form hosts. Scale bars A, B = 100 nm, D = 1 µm. **\*^Δ^**Illustrated zoom parts.

We observed cells from different types infected with Shield VLPs: amoeboid forms 3 µm in diameter and different types of flagellated microeukaryotes with a head ranging from 1.8 µm to 6 µm with one or two flagella (**Fig. 5D, Extended data Fig. 3B**). The nature of their hosts and their auto- and/or heterotrophic trophic mode remain to be determined. However, some of them did not exhibit or lacked pigment content under fluorescence microscopy. This putative broad host range was also suggested by positive correlations between Shield VLPs and almost all the microbial compartments mentioned above (p-value < 0.01). We assumed that Shield VLPs infect flagellated heterotroph microeukaryotes. Indeed, very rare large viruses such as *Cafeteria roenbergensis* virus^58^ and Klosneuviruses^54,59^, which are capable of infecting them, have been morphologically described^60^, and no ecological description (*in situ* temporal or spatial dynamic) is available. Shield VLPs are ubiquitous and were encountered in all three lakes studied here. We quantified Shield VLPs with remarkable abundances reaching 13.7 x 10^6^ VLPs.mL^-1^ and with a dynamic characterized by irregular phases of development (**Fig. 5C**).

With an observed mean BS of 27 (**Fig. 5D, Extended data Fig. 3B**), we estimated Shield VLPs to be involved in the mortality of 1.3 x 10^7^ hosts.L^-1^.day^-1^ (**Table 1**).

The ubiquitous distribution of Shield VLPs, which developed at different times of the year under contrasting environmental conditions, was reinforced by our observations of numerous Shield morphotypes in marine systems in which they can reach 8.3 x 10^4^ VLPs.mL^-1^ in surface area (**Extended data Fig. 3C**) as well as by their previous observation in soils^26^. Coupled with the pleomorphism of Shield VLPs and its potential host range, we suggest that this could represent a new generalist phylum, not considered until now, which plays a major ecological role in the regulation of microeukaryotes in many environments.

Finally, we also recorded striking rare forms of Shield-like VLPs (less than 1% of our observations) bearing tails or fibrils or in tandem (**Extended data Fig. 4**), up to 600 nm in diameter. We cannot certify the presence of these tail forms in the cytometric gate corresponding to the Shield VLPs.

#### Giant icosahedral virus-like particles: Novelties in common large viruses

This population, sorted by FC and verified by TEM, corresponded to giant icosahedral VLPs (GIV) (**Fig. 6A**). Like Ham and Shield VLPs, their cytometric signature suggested a large genome. A particular feature was their high signal on fluorescence associated with SSC, suggesting high internal complexity (**Fig. 1A, 6A**).

**Fig. 6.**
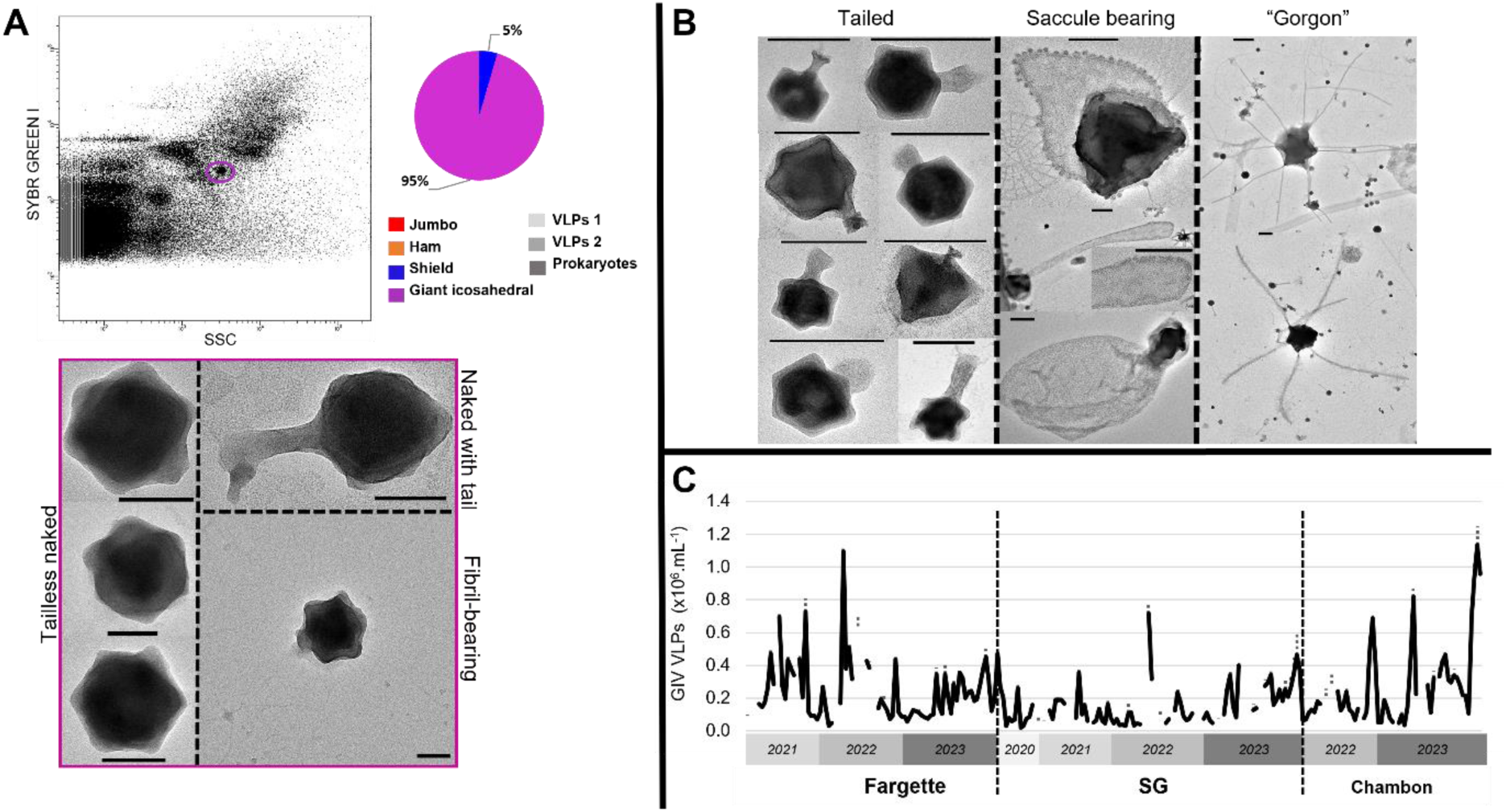
Detection, morphological and ecological characterization of giant icosahedral virus-like particles (GIV) detected in eutrophic French lakes. **A**, Flow cytometry (FC) detection of remarkable population of GIV and diversity of entities recorded by transmission electron microscopy after FC sorting in the corresponding population with micrographs of the GIV sorted. **B**, Negative staining electron micrographs of various GIV, tailed, bearing a saccule or “Gorgon”. Scale bars A = 100 nm, B = 200 nm. **C,** Seasonal dynamics of the abundance of GIV, in lakes Fargette, SG and Chambon. Each data represents the average of triplicates, dotted lines indicate standard deviation. n=209.

In this population, three morphotypes were observed: (i) tailless naked giant icosahedron (79% of the population in Chambon on January 10, 2024) of 212 nm in diameter; (ii) tail giant icosahedron (20%) of 180 nm in diameter with a tail of 150 nm length; and (iii) fibrils bearing giant icosahedron (< 1%) of 230 nm in diameter (**Fig. 6A**). The morphologies and sizes of the first two groups are similar to those observed for members of the *Megaviricetes* class, whereas no affiliation was proposed for the observed tailed types. Illustrations of various tailed GIV are provided in **Fig. 6B**. Finally, we also reported rare (less than 1% of the GIV observations) and GIV with a saccule (up to 446 nm icosahedral diameter) and morphotypes with tubular appendages (**Fig. 6B**) previously recorded in forest soils as “Gorgon”^26^. We cannot certify the presence of these amazing tailed forms in the cytometric gate corresponding to the GIV.

Abundances of GIV could reach 1.1 x 10^6^ VLPs.mL^-1^ and showed a dynamic with an irregular phase of development (**Fig. 6C**). GIV VLPs were recorded year-round, suggesting that they were dominant with numerous phylotypes and had a potential broad host range. This suggestion was also supported by the correlations between GIV and almost all the microbial compartments (p-value < 0.01).

Our data are among the first to document the absolute abundance of this presumably diverse population of *Megaviricetes*. The limited data available to date stemmed from specific isolated and fully described marine phytoplankton viruses during algal bloom^49–51,61,62^.

In the absence of a clear and visible burst event, we used a theoretical BS of 50 to estimate the maximum mortality of 3 x 10^6^ hosts.L^-1^.day^-1^ (**Table 1A**).

#### Snake and Sword virus-like particles: New dynamic and specific viruses of microeukaryotes

We report on the discovery of two VLPs above 200 nm, which we called Snake and Sword VLPs on account of their shape (**Fig. 7-8**). One particularity was that they are undetectable by FC based on the measured parameters. This lack of FC detection using dsDNA dye raises the question about the nature of their nucleic acid, probably non-dsDNA. Another explanation is that the signal is hidden within the bulk of VLPs1, VLPs2, or prokaryotes. However, despite their high abundances during the infection period, we found no correlation between TEM counts and any viral cytometric population.

**Fig. 7.**
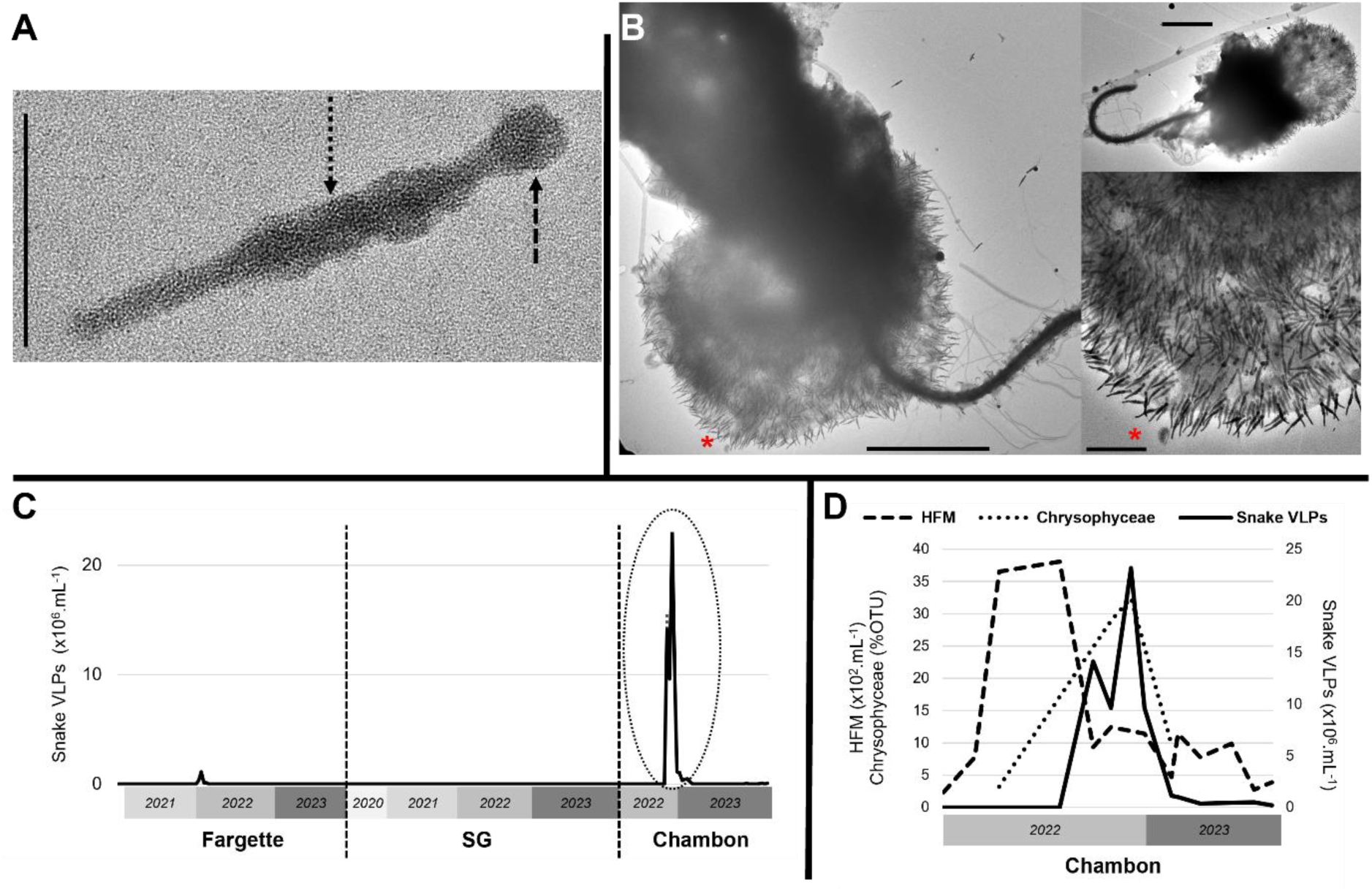
Detection, morphological and ecological characterization of Snake virus-like particles (VLPs) detected in eutrophic French lakes. **A**, Negative staining electron micrographs of Snake VLP showing two prominent parts (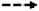) and (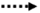). **B**, Negative staining electron micrographs of Snake VLPs derived from a lytic event in a microeukaryote host. **\*** Illustrated zoom part. **C**, Seasonal dynamics of the abundance of Snake VLPs, in lakes Fargette, SG and Chambon. Each data represents the average of triplicates, dotted lines indicate standard deviation. n=252. Scale bars A = 200 nm, B = 2 µm with scale in zoom part = 500 nm. **D,** Focus on remarkable infection periods on the dynamics of Snake VLPs, heterotrophic flagellated microeukaryotes (HFM) and relative abundance of Chrysophyceae class (% OTU) in lake Chambon from September 23, 2022 to February 17, 2023.

**Fig. 8.**
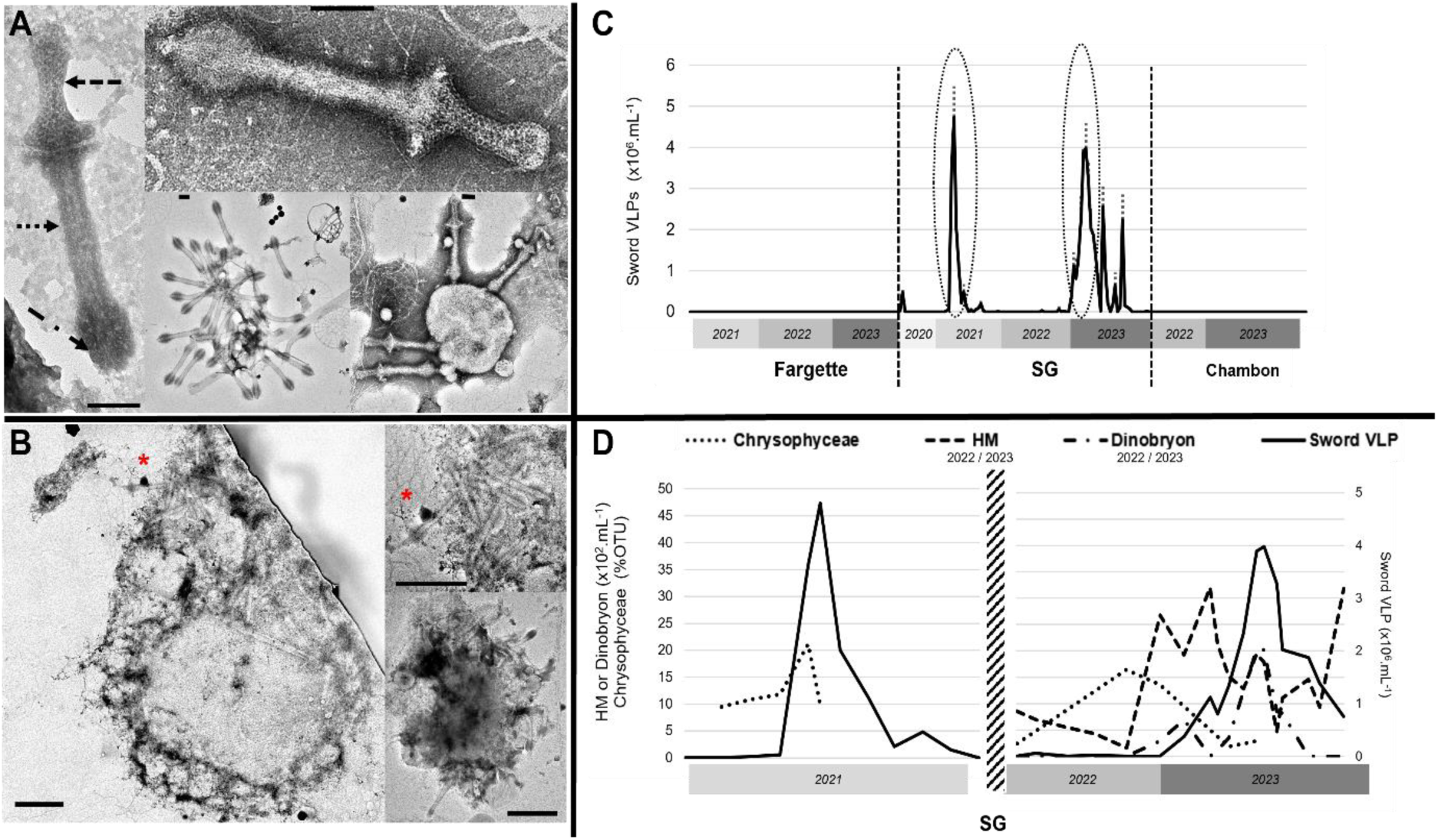
Detection, morphological and ecological characterization of Sword virus-like particles (VLPs) detected in eutrophic French lakes. **A**, Negative staining electron micrographs of Sword VLPs with a head ()-tail () morphology terminated by fibers (). **B**, Negative staining electron micrographs of Sword VLPs derived from a lytic event in a microeukaryote host. **\*** Illustrated zoom part. Scale bars A = 100 nm, B = 500 nm. **C,** Seasonal dynamics of the abundance of Sword VLPs, in lakes Fargette, SG and Chambon. Each data represents the average of triplicates, dotted lines indicate standard deviation. n=252. **D**, Focus on remarkable infection periods on the covariations of sword VLPs, heterotrophic microeukaryotes (HM) and Dinobryon sp., and relative abundance of Chrysophyceae classes (% OTU) in lake SG from February 04, 2021 to June 30, 2021 and from October 18, 2022 to April 12, 2023.

Snake VLPs are elongated with a length of 226 nm (**Fig. 7A**). Their neck delimits a spherical part measuring 30 nm in diameter with a cylindrical part of 30 nm in diameter and 100 nm in length and ends with a thinner cylindrical part measuring 15 nm in diameter and 78 nm in length. They have a polarity with a preferential orientation, with the spherical part pointing toward the host (**Fig. 7B**). Snake VLPs infect a flagellated microeukaryote 5 µm in diameter, identified as a heterotrophic Chrysophyceae on morphological and light microscopy criterion^63^ (**Fig. 7B**). These observations were corroborated by the simultaneous dynamics of the abundance of Snake VLPs and Chrysophyceae approximated by 18S metabarcoding data, which were particularly noticeable as Chrysophyceae-dominated eukaryote communities at the time of the Snake VLP bloom (**Fig. 7D**). Snake VLPs were predominant in Lake Chambon, with a maximum abundance of 23.2 x 10^6^ VLPs.mL^-1^. Their dynamic was characterized by a sudden and massive development strategy with phases of infection alternating with long phases without detection (**Fig. 7C**). The infectivity of Snake VLP was high, with a production capacity exceeding 1,200 VLPs per lysed host. At peak infection, all host cells observed in TEM were infected.

Sword VLPs are a head-tail-fiber VLP measuring 635 nm in length (**Fig. 8A**). This viral type is characterized by an elongated dumbbell-shaped head measuring 220 nm in length and between 45 and 135 nm in width, with a sheath-like tail of 370 nm in length and 60 nm in width and fibers distributed at the end of the tail. A thin 9.5 nm thick groove separates the presumed head from the tail. The head has a distinctive substructure resulting from a patterned arrangement of 6 nm diameter subunits. Morphologically, the closest relative appears to be a filamentous VLP previously imaged in a freshwater lake^64^. We reported our observation of infected burst unicellular microeukaryotes of at least 3-4 µm in diameter, but unfortunately, we cannot specify their identity (**Fig. 8B**). Metabarcoding data showed that at time of the VLP bloom, eukaryote communities were dominated by Chrysophyceae (**Fig. 8D**). We found Sword VLP only in Lake SG. Their dynamic, reaching 4.7 x 10^6^ VLPs.mL^-1^, was characterized by a sudden and massive development strategy (**Fig. 8C**). The BS observed for Sword VLPs appeared to be around 15.

On an expanded spatiotemporal scale, we were unable to identify significant correlations (p-value > 0.01) between Snake or Sword VLPs and the autotrophic microbial compartments considered in this study. Nevertheless, focusing on the remarkable viral infection peaks from September 23, 2022, to February 17, 2023, at Lake Chambon and from October 18, 2022, to April 12, 2023, at Lake SG, we showed a clear interaction in the “prey-predator” model between Snake VLPs and heterotrophic flagellated microeukaryotes (**Fig. 7D**) and between Sword VLPs and heterotrophic microeukaryotes or Chrysophyceae, respectively (**Fig. 8D**).

The interaction between heterotrophic microeukaryotes, Chrysophyceae, and Sword VLPs was also supported by various observations of numerous Sword VLPs inside a heterotrophic microeukaryotes associated with a putative Chrysophyceae lorica (probably Dinobryon sp.) and numerous Sword VLPs around a lorica remnant (**Extended data Fig. 5**).

These dynamic analyses support the TEM observations and metabarcoding data showing a putative heterotrophic microeukaryote host for Snake VLPs. The trophic status of the Sword VLP host remains uncertain. Precise identification will require host isolation.

Finally, the reproducibility of morphological observations of the Snake and Sword VLPs and their host coupled with their development strategy suggest that they are highly specialist VLPs.

These results showed the high involvement of Snake and Sword VLPs in the demise of their host population with a maximum mortality of 0.1 and 2.4 x 10^7^ hosts.L^-1^.day^-1^ for Snake and Sword VLPs, respectively (**Table 1A**).

#### Ecological significance of large microeukaryote virus-like particles

This study represents one of the first time series of large microeukaryote VLPs and provides novel insights into their numerical significance in aquatic systems. Detection limitations such as the inability to detect Snake and Sword VLPs using FC will likely lead to an underestimation of the total number of large microeukaryote VLPs. Nevertheless, our data emphasize the numerical importance of these entities in freshwater systems, with total abundances ranging from 0.2 up to 28.4 x 10^6^ VLPs.mL^-1^. As such, large microeukaryote VLPs accounted for between 0.1 and 2.7%, 0.1 and 6.8%, and 0.3 and 10.8% of the total viral stock in Lakes Fargette, SG, and Chambon, respectively (**Extended data Fig. 6 and Table 1B**). These variable contributions point to the alternating control of microbial systems by large microeukaryote VLPs versus smaller viruses (mostly phages). Of notable interest, this study also reports the significant contribution of large VLPs in an offshore marine site and particularly Shield VLPs, thus expanding the distribution of this morphotype to all aquatic systems.

On an annual basis, the productive periods of large VLPs occurred mainly in spring, with some exceptions such as the autumn peak in 2022 in Lake Chambon associated with Snake VLPs and the summer peak in 2022 in Lake Fargette. This strong temporal dynamism was also underlined by the rapid and unexpected successions of specific large VLPs. Some morphotypes (Ham, Shield, Snake, Sword) can evolve from undetectable levels to over 80% of the total large VLPs in a matter of days. At other times, a mixed community of large VLPs was observed in equal proportions (**Extended data Fig. 6A)**. Our results were consistent with the “Bank model”^65^, which suggests that only a small fraction of the viral community is active and abundant at any given time, while most populations are rare and dormant, forming a seed bank that can “Kill-the-Winner” when hosts reach critical abundance thresholds^66^. It is worth noting that naked GIV were only dominant once in Lake SG (55% of the total large VLP community on May 30, 2022) and that they represent only 12% of all the combined data on average. Thus, large VLPs not previously considered morphologically (Ham, Shield, Snake, Sword) could be prevalent in the microeukaryote ecology of the aquatic environment.

To decipher the effect of large microeukaryote VLPs on host communities, we monitored the dynamics of their total abundance concomitantly with those of microbial communities (abundance and diversity). Over the entire study period, the total abundance of large VLPs correlates with the total microeukaryotes (R² = 0.4, p-value < 0.02). Alongside the observations of “prey-predator” patterns between autotrophic microeukaryotes and Ham VLPs and between heterotrophic microeukaryotes and Snake VLPs (**Fig. 4D, 7D)**, these findings suggest that both autotrophic and heterotrophic microeukaryotes contributed significantly and variably to the stock of large VLPs. We observed no significant correlation with abiotic parameters (**Extended data Fig. 7**), thus confirming the importance of the microbial environment in the control of large VLPs. The high values of maximum ratios between the total large microeukaryote VLPs and microeukaryotes (325, 949, and 1463 in Lakes Fargette, SG, and Chambon, respectively) (**Table 1B**) point to the significant potential of large VLPs to control microeukaryote populations.

We estimated the removal of NMP by large microeukaryote VLPs by calculating the ratio between the theoretical viral-induced mortality of microeukaryotes and net microeukaryote production. NMP removal does not account for viral decay or microeukaryote dynamics during the measurement period, which was limited to a maximum of 2 weeks. It should also be noted that small microeukaryotes (i.e., nanoeukaryotes) are difficult to distinguish from large prokaryotes or fungi spore by light microscopy or FC, and are probably underestimated in microeukaryote counts, which can lead to an overestimation of this ratio. The values ranged from -0.7 to 0.2 in Lake Fargette, from -10 to 37 in Lake SG, and from -6 to 15.6 in Lake Chambon (**Table 1B**). This significant variation indicates that at specific times, large microeukaryote VLPs can control the stock of microeukaryotes and contribute to the observed abundance dynamics (**Extended data Fig. 8**).

The potential influence of large VLPs on microeukaryote succession was assessed by analyzing the temporal dynamics of microeukaryotic diversity (Shannon index) in relation to the abundance of large VLPs of microeukaryotes (**Fig. 9**). Our findings showed that episodes of viral production, regardless of the dominant viral type, coincided with a decrease in eukaryote diversity. This pattern is particularly evident in Lake Chambon during the peak of Snake VLPs, which are suspected to control a population of heterotrophic flagellates that prey on other microorganisms^67^. During the massive proliferation of heterotrophic flagellates, which likely causes a drop in diversity, Snake VLPs not only regulate their abundances but also release their predation pressure, which could in turn free up niches for the proliferation of other microeukaryotes. The effects of large VLPs may also be cumulative as in the example of the simultaneous development of Snake VLPs and GIV in Lake Chambon, or they may follow each other successively as in the example of the successive production of Ham and Shield VLPs in Lake Fargette. Although it is clear that large VLP communities were an essential factor in regulating the succession of microeukaryote communities in our study sites, the lack of information regarding their life traits (e.g., infection mode, host specificities) or subcellular interactions limits our understanding of their overall impact. Dedicated efforts to isolate and characterize the interactions of large VLPs must be pursued in order to gain insights into the role of this underexplored, yet significant component of the virosphere by means of cultures, subcellular observations, and transcriptomic analyses^3,25,68^.

**Fig. 9.**
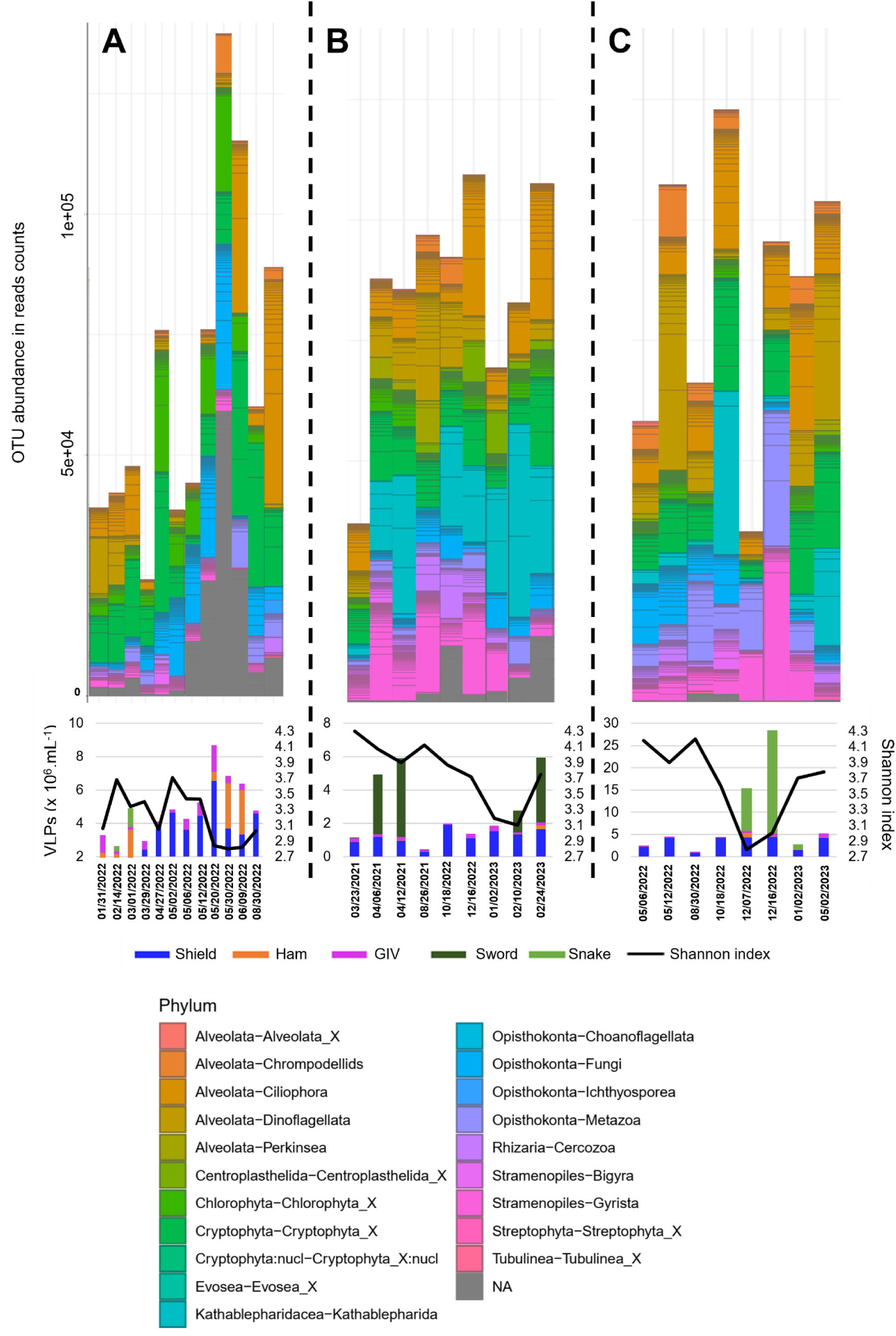
Temporal dynamics of eukaryote OTU abundance in reads counts, and of eukaryote diversity index (Shannon index) compare to viral abundance of Ham, Shield, Snake, Sword and giant icosahedral viruses (GIV) in lakes Fargette **(A)**, SG **(B)** and Chambon **(C)**.

Surprisingly, the minimum values for the abundance, microeukaryotic virus/total virus ratio, microeukaryote VLP and microeukaryote ratio, and NMP removal by large microeukaryotic VLPs were encountered in Lake Fargette, which presents the highest microbial abundances and pigment contents (**Table 1, Extended Data** Fig. 6 and 8), thus suggesting better control in less eutrophic environments.

### Conclusions and perspectives

At a large temporal scale, the combination of FC and TEM led to original and unexpected advances in determining the diversity and ecology of large aquatic VLPs. Indeed, we shed light on the astonishing phenotypic diversity of large VLPs in aquatic ecosystems, with the identification of specific cytometric populations and four novel morphotypes never previously described. These viruses, which infect prokaryotes or microeukaryotes undoubtedly play a role in the control of their host population. Each type of VLP probably represents different taxonomic levels and specializations (from specialist to generalist). Some notably seem to infect heterotrophic microeukaryotes, for which few viruses have been reported. The global community of large VLPs, considered here for the first time on a large temporal scale, showed marked dynamics in terms of both their abundance and diversity, with each of the studied types either succeeding or accumulating. We suggest that these viral communities have a very strong impact on ecosystem functioning by controlling the dynamics of prokaryotic and eukaryotic communities, both autotrophic and heterotrophic, and by controlling their diversity, which implies their major role, hitherto unimagined at such a scale, in the functioning of the ecosystem.

These discoveries significantly impact our perception of the virosphere, its diversity, and the role played by large viruses in the dynamism of their hosts and the functioning of aquatic food webs. This study urges us to investigate the genomic characterization of these VLPs and to identify their host to fully explore these concepts. Advances in metatranscriptomic technologies will allow us to better clarify their role in the matter and energy flow of ecosystems by gaining access to their cellular and subcellular levels.

## Data availability

Relevant data supporting the key findings of this study are available in the article and the Supplementary Information file. All raw data generated du the current study are available from the corresponding author upon request.

## Supporting information

Supplemental files Billard et al. 2024

## Acknowledgements

This project received financial support from the CNRS through the MITI interdisciplinary programs (X-life 2018-2019, Origines 2020-2021, and PIB 2023-2024). We also benefited from the following: (i) the CPER 2015–2020 SYMBIOSE challenge program (French Ministry of Research, UCA, CNRS, INRA, Auvergne-Rhone-Alpes Region, FEDER), which supported the PhD fellowship of M.F.; (ii) funding from the French government IDEX-ISITE initiative 16- IDEX- 0001 (CAP 20–25); and (iii) the contribution of the APERO project funded by the National Research Agency under the grant APERO [grant number ANR ANR-21-CE01-0027] and by the French LEFE-Cyber program (CNRS, INSU). We would also like to thank the CYSTEM–UCA PARTNER Platform (Clermont-Ferrand, FRANCE) for their technical support and expertise.

We are grateful to Dr Mart Krupovic for his expertise and numerous discussions and for his help in drafting the manuscript.

We are indebted to all those who contributed to these efforts.

## Author information

Authors and Affiliations

**Laboratoire Microorganismes : Génome et Environnement (LMGE), UMR CNRS 6023, Université Clermont-Auvergne, F-63000 Clermont-Ferrand, France**

Hermine Billard, Maxime Fuster, François Enault, Jean-François Carrias, Léa Fargette, Margot Carrouée, Perrine Desmares, Télesphore Sime-Ngando, Jonathan Colombet

**Génomique Métabolique, Genoscope, Institut François Jacob, CEA, CNRS, Univ. Evry, Université Paris-Saclay, Evry, France**

Tom O. Delmont

Sorbonne Université, CNRS, Station Biologique de Roscoff, FR2424, Roscoff, France

Gwenn Tanguy

Sorbonne Université, CNRS, Station Biologique de Roscoff, UMR 7144, Roscoff, France

Estelle Bigeard, Pauline Nogaret, Anne-Claire Baudoux

UMR CNRS 8187 LOG, Université Littoral Côte d’Opale, Université de Lille, Wimereux, France

Urania Christaki

## Contributions

J.C. supervised this research. H.B. and J.C. conceived the project, designed and led the experiment. E.B., G.T., P.N., A.C.B., and U.C. provided the marine samples. H.B. and J.C. co- wrote the manuscript. J.F.C., H.B., and P.D. analyzed the diversity of the microeukaryotes. All authors contributed to the data analysis and discussion.

## Ethical declarations

Competing interests

The authors declare no competing interests.

