## Supplemental files Billard et al. 2024 for "Unexpected diversity and ecological significance of uncultivable large virus-like particles in aquatic environments"

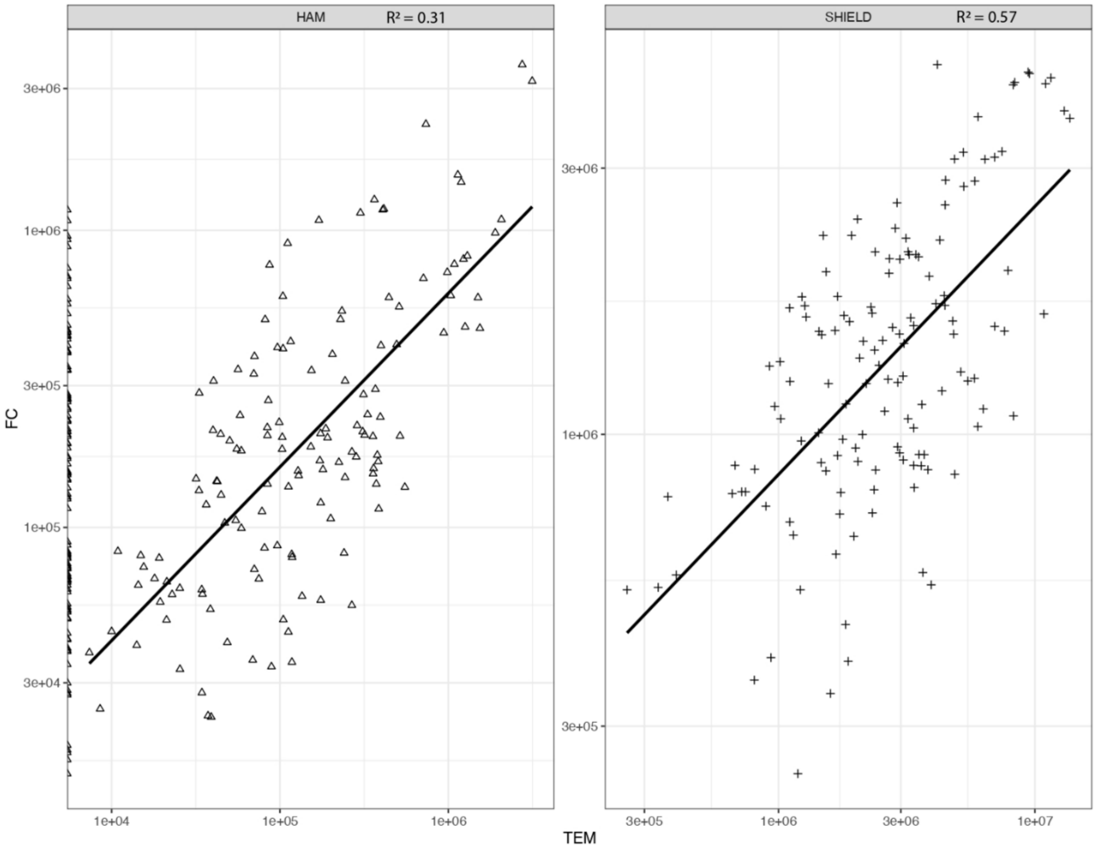


**Extended Data Fig. 1** Spearman correlative analysis (all data from three lakes combined) between counts of Shield or Ham virus-like particles by flow cytometry (FC) compared to counts by transmission electron microscopy (TEM). p values < 0.001. n Ham = 247, n Shield = 135.

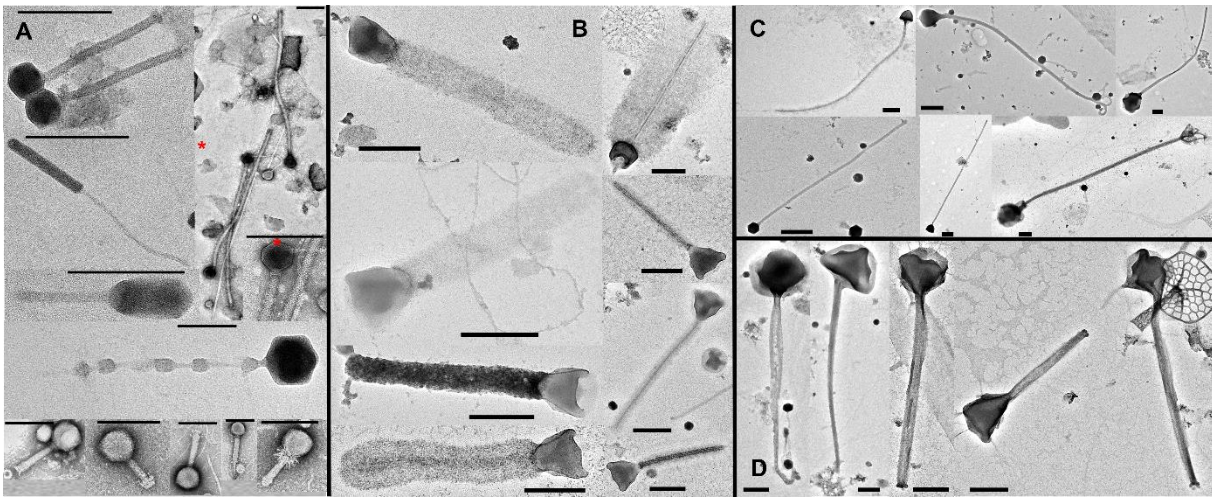


**Extended Data Fig. 2** Negative staining electron micrographs of tailed virus-like particles (VLPs) detected in eutrophic French lakes. **A,** Jumbo-like phages. **B,** sheath tailed VLPs. **C,** naked elongated VLPs. **D,** tubular tailed VLPs. Scale bars = 200 nm.

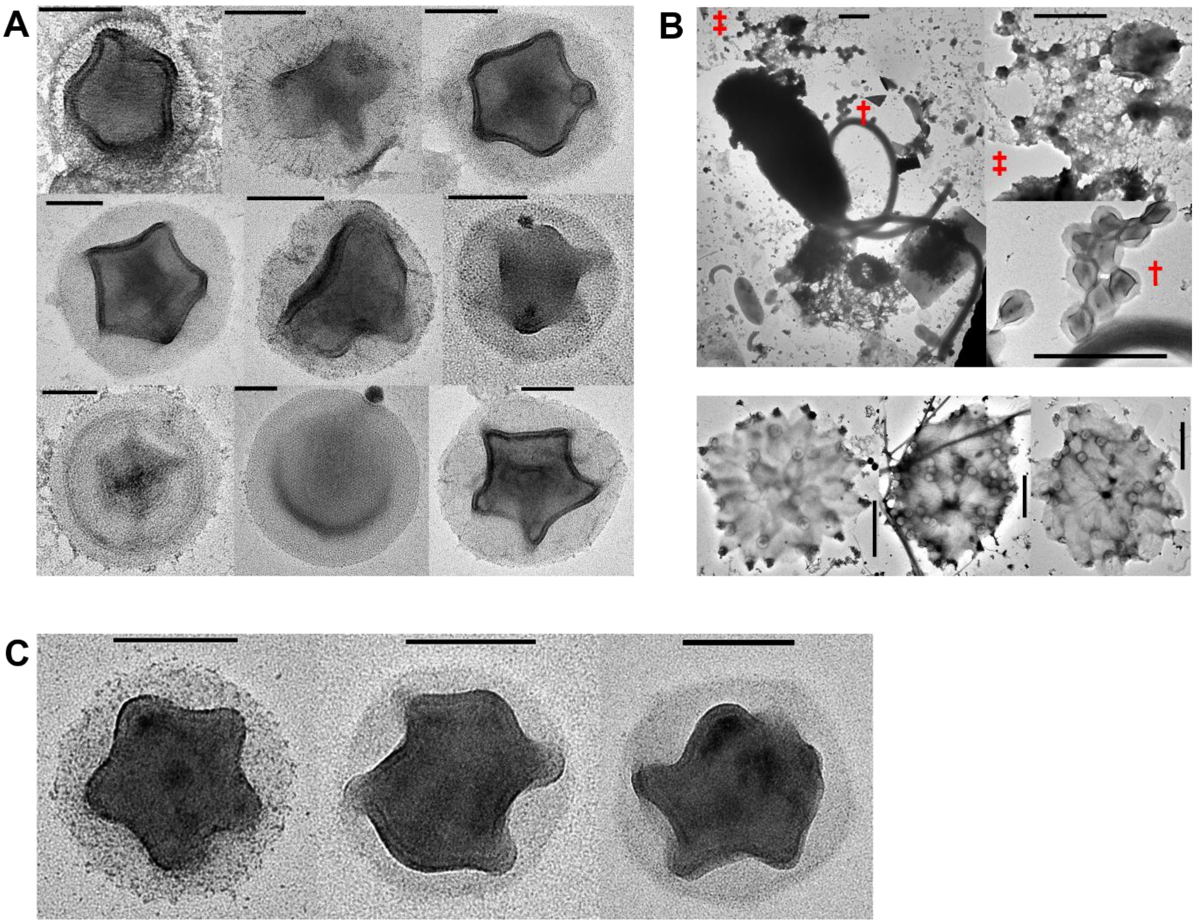


**Extended Data Fig. 3 A**, Negative staining electron micrographs of various Shield virus-like particles (VLPs) Scale bars= 100 nm. **B**, Negative staining electron micrographs of Shield VLPs derived from lytic events of flagellated microeukaryote or amoeboid form hosts. Scale bars = 1µm. ‡† Illustrated zoom parts. **C**, Negative staining electron micrographs of Shield VLPs detected in marine environment. Scale bars = 100 nm.

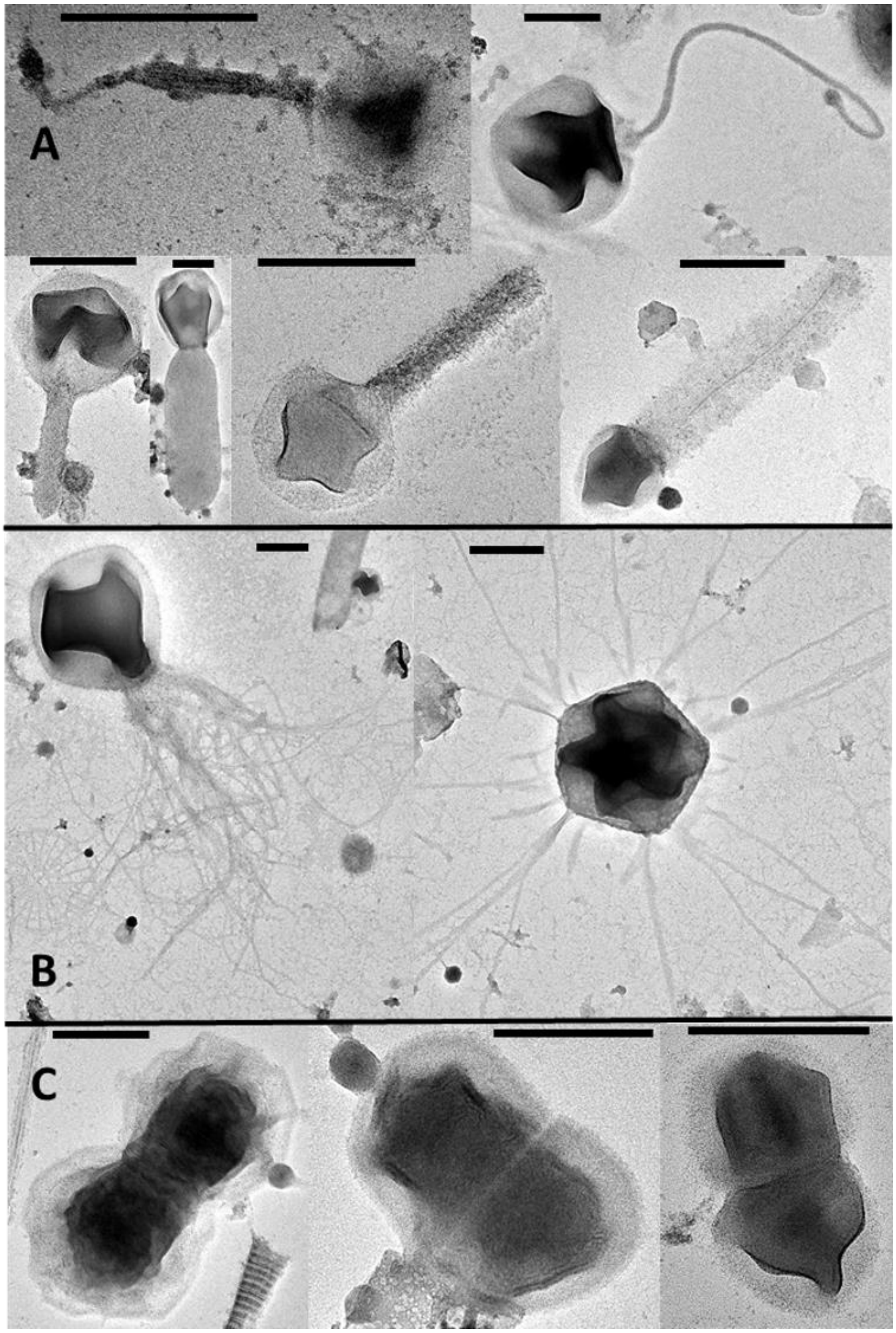


**Extended Data Fig. 4** Negative staining electron micrographs of Shield virus-like particles (VLPs) detected in eutrophic French lakes. **A**, Tailed Shield VLPs. **B**, Fibrillar Shield VLPs. **C**, Tandem Shield VLPs. Scale bars = 200 nm.

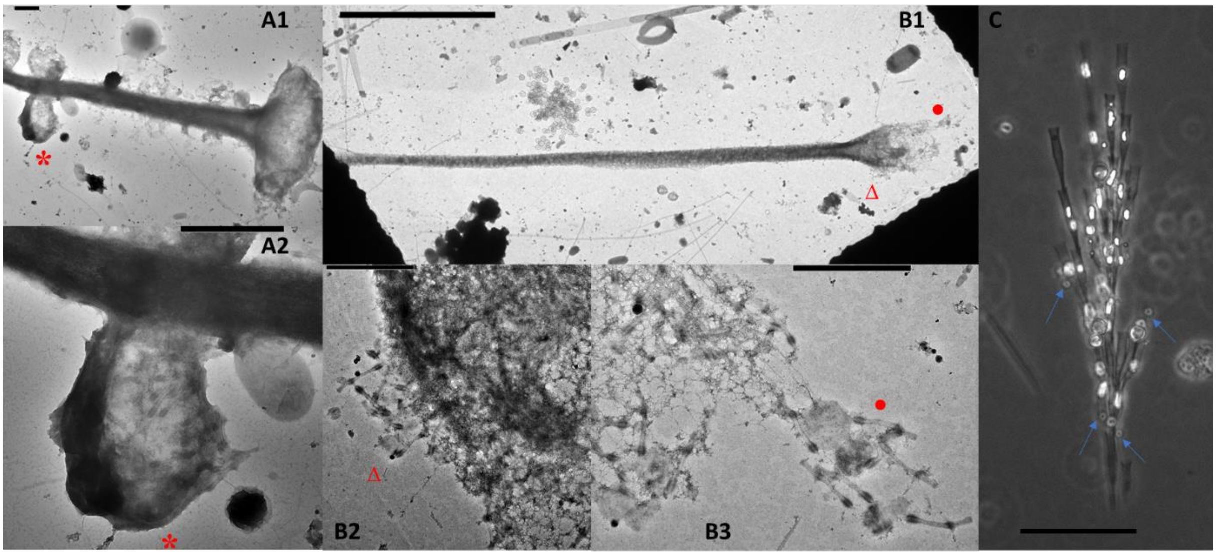


**Extended Data Fig. 5** Putative interactions between Sword virus-like particles (VLPs), heterotrophic microeukaryotes (HMs) and Chrysophyceae, probably Dinobryon sp. **A1** and zoom part **A2**, negative staining electron micrographs of Sword VLPs detected in an HM associated with a putative Dinobryon sp lorica. **B1** and zoom parts **B2** and **B3**, negative staining electron micrographs of Sword VLPs detected around a putative Dinobryon sp. lorica after the burst of its host. **C**, Light image of a Dinobryon sp. colonized by HM (arrows). *Δ• Illustrated zoom parts. Scale bars A1, A2, B2, B3 = 1 µm; B1=10 µm; C = 100 µm.

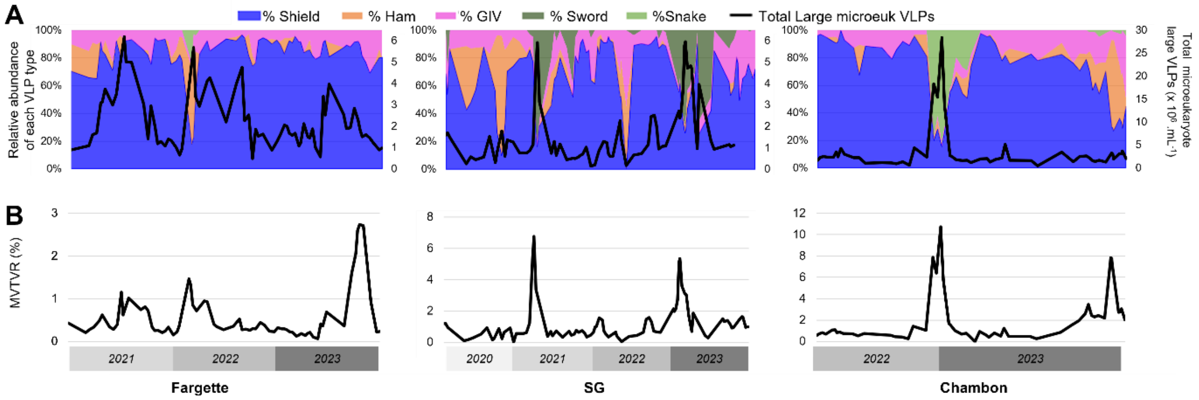


**Extended data Fig. 6** Seasonal dynamics in lakes Fargette, SG and Chambon (n = 71, 81, 53 respectively) of relative abundances of Shield, Sword, Ham, Snake, GIV and total microeukaryote large virus-like particles (VLPs) abundance **(A)** and of the large microeukaryote VLPs to total virus ratio (MVTVR) **(B)**. Each data was obtained from the average of triplicates. Only data for which counts of all viruses-like particles considered are available are displayed.

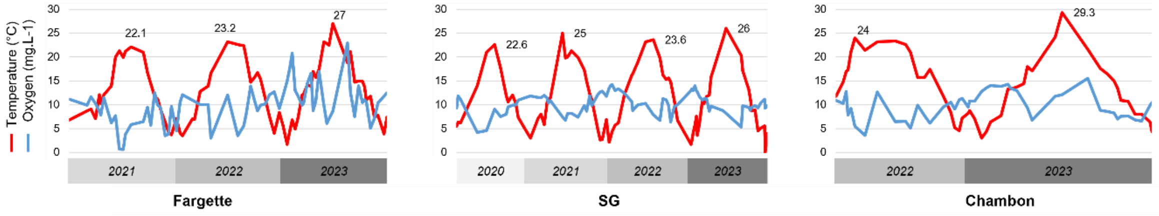


**Extended Data Fig. 7** Seasonal dynamics of temperature and oxygen contents in lakes Fargette, SG and Chambon (n = 71, 81, 53 respectively). Only data for which counts of all viruses-like particles considered are available are displayed.

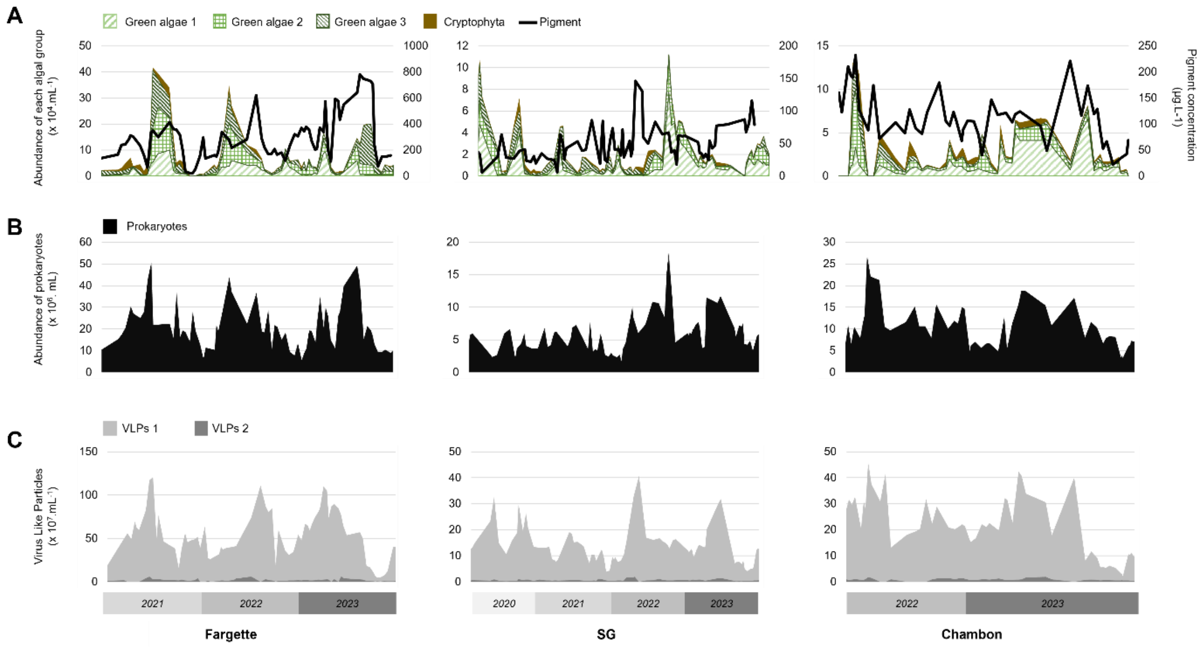


**Extended Data Fig. 8** Seasonal dynamics of autotrophic microeukaryotes abundance and total pigment concentration (**A**), prokaryotes abundance (**B**), and VLPs 1 and 2 respectively abundances (**C**) in lakes Fargette, SG and Chambon (n = 71, 81, 53 respectively). Each data, excepted autotrophic microeukaryotes, represent the average of triplicates. Only data for which counts of all viruses-like particles considered are available are displayed.
